# A Small Subset of Cytosolic dsRNAs Must Be Edited by ADAR1 to Evade MDA5-Mediated Autoimmunity

**DOI:** 10.1101/2022.08.29.505707

**Authors:** Tao Sun, Qin Li, Jonathan M. Geisinger, Shi-Bin Hu, Boming Fan, Shichen Su, Waitang Tsui, Hongchao Guo, Jinbiao Ma, Jin Billy Li

## Abstract

The innate immune system detects viral infection via pattern recognition receptors and induces defense reactions such as production of type I interferon^1^. One such receptor, MDA5, is activated upon the recognition of double-stranded RNAs (dsRNAs) that are often produced during viral replication^2^. Endogenous dsRNAs evade MDA5 activation through RNA editing by ADAR1, thus preventing autoimmunity^3-5^. Among the large number of endogenous dsRNAs, the key substrates whose editing is critical to evade MDA5 activation (termed as immunogenic dsRNAs) remain elusive. Here we reveal the identity of human immunogenic dsRNAs, a surprisingly small fraction of all cellular dsRNAs, to fill the gap in the ADAR1-dsRNA-MDA5 axis. We found that, in contrast to previous findings^6,7^, the immunogenic dsRNAs were highly enriched in mRNAs and depleted of introns, an expected indication of *bona fide* substrates of cytosolic MDA5. The immunogenic dsRNAs, in contrast to non-immunogenic dsRNAs, tended to have shorter loop between the stems, which may facilitate dsRNA formation. They also tended to be enriched at the GWAS signals of common inflammatory diseases, implying that they are truly immunogenic. We validated the MDA5-dependent immunogenicity of the dsRNAs, which was dampened following ADAR1-mediated RNA editing. We anticipate that a focused analysis of immunogenic dsRNAs will greatly facilitate the understanding and treatment of cancer and inflammatory diseases in which the important roles of dsRNA editing and sensing continue to be revealed^8-13^.

## Main

Adenosine-to-inosine (A-to-I) RNA editing, which is catalyzed by adenosine deaminase acting on RNA (ADAR) enzymes, is one of the most common RNA modifications in metazoan^14-16^. ADAR binds to double-stranded RNAs (dsRNAs) and converts specific adenosines to inosines, which are recognized as guanosines by the cellular machinery^17^. RNA editing sites are predominantly located in repetitive sequences that have a neighboring, inverted-orientated repeat in the genome, thus forming long dsRNAs with multiple adenosines edited^18,19^. In humans, millions of editing sites have been documented^20-23^, the vast majority of which are located in the primate-specific *Alu* repeats. Similarly, in mice, the vast majority of editing events are in the *Alu*-like SINE elements (e.g., B1, B2). Following the primate-rodent split during evolution, *Alu* and B1/B2 elements spread independently in each of the two genomes in a seemingly random manner, suggesting species-specific pools of long dsRNAs formed by inverted repeats.

In mammals, there are two enzymatically active ADAR enzymes, ADAR1 and ADAR2, and ADAR1 is expressed as two isoforms, p110 in the nucleus and p150 in the cytoplasm^24-29^. *Adar2*^*-/-*^ mice die within 3 weeks after birth with seizure, primarily due to the lack of editing at a single recoding site in *Gria2*^30^. *Adar1*^*-/-*^, *Adar1p150*^*-/-*^ (lacking the cytoplasmic ADAR1 isoform) and catalytically inactive ADAR1 (*Adar1*^*E861A/E861A*^) mice are embryonic lethal with massive interferon (IFN) activation^4,31-33^. However, the key editing substrate(s) of ADAR1 have been elusive to date.

The biological function of ADAR1 RNA editing has been unveiled in recent years. The catalytically inactive ADAR1 (*Adar1*^*E861A/E861A*^) mouse, can be rescued to full life span by the ablation of MDA5, a host dsRNA sensor in the innate immune system^4^. Similarly, the embryonic lethality of *Adar1*^*-/-*^ mice was rescued by removal of MDA5 or its downstream adaptor protein MAVS^3,5^. MDA5, encoded by *IFIH1*, is a Pattern Recognition Receptor (PRR)^1^ that recognizes long dsRNA often produced through viral replication^2^. Many copies of MDA5 form a filament along the length of dsRNA, which activates the downstream signaling cascade involving MAVS, TBK1, and IRF3 to induce the expression of type I IFN and eventually interferon-stimulated genes (ISGs)^34^. In the absence of ADAR1 editing, endogenous “self” dsRNAs would be recognized by MDA5 as “non-self” to trigger deleterious interferon responses. Therefore, the key physiological function of ADAR1 mediated RNA editing is to suppress MDA5-mediated dsRNA sensing.

The established ADAR1-dsRNA-MDA5 axis is consistent with human genetics. Loss-of-function (LOF) mutations of ADAR1 have been found in rare autoimmune diseases such as Aicardi-Goutières Syndrome (AGS) that mainly affects the brain and clinically mimics in-utero-acquired viral infection despite the absence of detectable virus^35,36^. AGS can also be caused by MDA5 gain-of-function (GOF) mutations that relax the dsRNA specificity^6,37^. Recently, we uncovered the role of the ADAR1-dsRNA-MDA5 axis in common autoimmune and inflammatory diseases^13^. Genetic variants associated with dsRNA editing levels are significantly enriched in GWAS signals for immune-related diseases. The genetic risk variants defined by GWAS collectively reduced editing levels of nearby dsRNAs, contributing to elevated type I IFN response and the subsequent increased disease risk observed in patients. In summary, human genetics data demonstrates the important roles of the ADAR1-dsRNA-MDA5 axis in immune-related diseases and calls for systematic identification of cellular dsRNAs with disease significance.

Of the large number of dsRNAs that are edited by ADAR1, which ones must be edited to avoid the MDA5 recognition and subsequent interferon responses? Such dsRNAs would be implicated as triggers in many inflammatory diseases. Hereafter, we define these key ADAR1 editing substrates as immunogenic dsRNAs, since they would lead to elevated immunogenicity unless they are edited. To identify immunogenic dsRNAs an intuitive approach is a direct capture of MDA5-bound dsRNA directly, by a method such as CLIP^38^. However, it is known that MDA5 promiscuously binds ssRNA and dsDNA, which does not result in downstream signal transduction^39^. Recently a more selective method was reported, in which total cytoplasmic RNA was incubated with recombinant MDA5 *in vitro*, and the RNAs bound by MDA5 filament (but not monomer) were protected from RNase digestion and later identified by RNA-seq^6^. While this study identified a large number of *Alu* repeats, the major weakness of this strategy is that the *in vitro* conditions may not recapitulate cellular or *in vivo* conditions. In addition, several lines of evidence suggest that there may be different repertoires of immunogenic dsRNAs in different cell types and tissues. For example, when *ADAR*^*-/-*^ human embryonic stem cells are differentiated into neural progenitor cells (NPCs) or hepatocyte-like cells (HLCs), interferon response and cell death phenotypes were only observed in NPCs, but not in HLCs^40^, consistent with the clinical manifestations of AGS patients with ADAR1 LOF mutations where inflammation is most pronounced in the brain^36^.

Here, we developed novel genetic and computational approaches to identify immunogenic dsRNAs in different cell types. Using HEK293T cells and NPCs, we identified both shared and cell type unique immunogenic dsRNA candidates. In both cell types, we found that only a very small subset of dsRNAs are immunogenic without being edited, in contrast to previous findings^6,7^. In addition to identifying *Alu* repeats, as expected, we discovered a novel class of dsRNAs that are formed by overlapping genes transcribed in opposite directions. We further characterized these immunogenic dsRNAs and identified features that distinguish them from non-immunogenic ones. Our work provides a framework to reveal immunogenic dsRNAs in the ADAR1-dsRNA-MDA5 axis, enabling future endeavors to identify key dsRNAs whose aberrant editing may lead to autoimmunity or immune-related diseases.

### RNA editing by ADAR1 suppresses MDA5-dependent interferon signaling in human cells

To model ADAR1 editing deficiency in human cells, we generated a deaminase inactive HEK293T cell line by mutating the glutamic acid 912 to alanine (E912A), resembling the E861A mutation in mice^4,41^ (**Extended Data Fig. 1a**). There are three ADAR1 alleles in the HEK293T genome. In the homozygous *ADAR1*^*E912A/E912A/E912A*^ (hereafter referred to as ADAR1^E912A^) cell line, RNA editing is almost entirely abolished and the residual amount of editing is presumably due to the lowly expressed ADAR2 (**Extended Data Fig. 1b–c**). In two heterozygous *ADAR1*^*+/E912A/E912A*^ and *ADAR1*^*+/+/E912A*^ lines we generated, we observed ADAR1 dosage-dependent levels of RNA editing (**Extended Data Fig. 1c–d**).

We observed no significant change in the expression of ISGs between ADAR1^E912A^ and WT (**Fig. 1a**). The lack of ISG induction is because endogenous MDA5 is not expressed in HEK293T cells, and exogenous expression of MDA5 leads to a drastic increase of ISG expression in ADAR1^E912A^ cells (**Fig. 1a**). In addition, we generated an *ADAR1*^*-/-/-*^ (hereafter ADAR1^KO^) mutant cell line (**Extended Data Fig. 1e–f**) and also observed MDA5-dependent ISG induction (**Extended Data Fig. 1g**), similar to a previous study^5^. To further confirm that the ISG induction in ADAR1^E912A^ cells is dependent on the MDA5 pathway, we knocked out MAVS, the gene that encodes the adaptor protein immediately downstream of MDA5. The ISG induction was completely abolished in the ADAR1^E912A^;MAVS^KO^ double mutant even in the presence of MDA5 (**Fig. 1b, Extended Data Fig. 1h**). Our results demonstrate that the lack of RNA editing by ADAR1 elicits the ISG induction via the MDA5-MAVS pathway in human cells, consistent with the *in vivo* mouse data^3-5^.

**Fig. 1:**
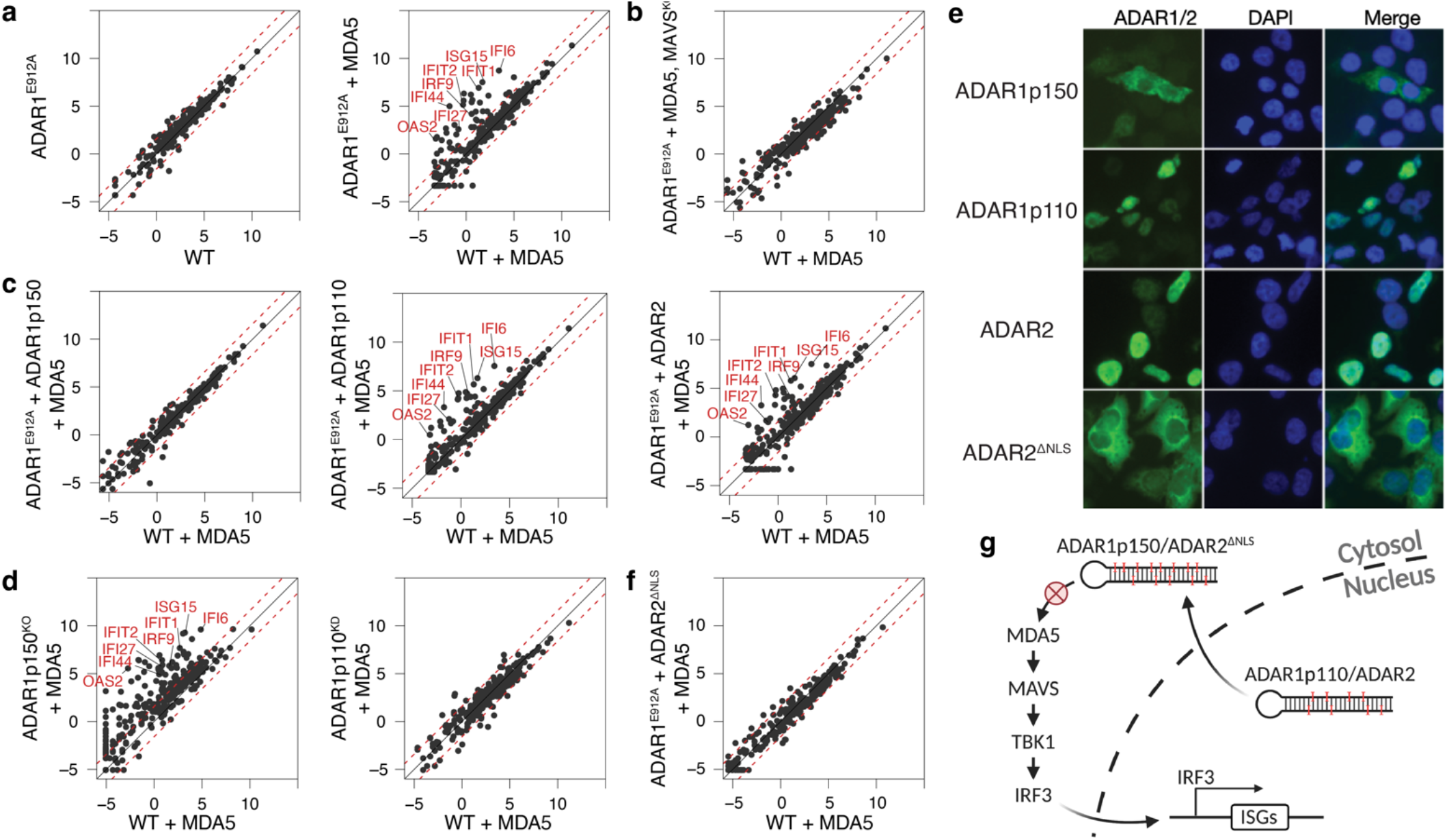
Cytosolic ADAR editing is essential to suppress MDA5-mediated dsRNA immunity. **a**, Expression comparison of ISGs (interferon stimulated genes) between WT and ADAR1^E912A^ cells without exogenous MDA5 (left) and with lentivirally expressed MDA5 (right). Threshold of three-fold change is marked in red dashes. A number of signature ISGs are labeled on the plot. The gene expression levels were measured by TPM (Transcripts Per Kilobase Million) and log2 transformed in this and other figures. **b**, Expression comparison of ISGs between ADAR1^E912A^ cells with exogenous MDA5 expressed and MAVS deleted and WT cells with exogenous MDA5 expressed. **c**, ADAR1p150 (left), ADAR1p110 (middle) and ADAR2 (right) were stably expressed in the ADAR1^E912A^ cells that have exogenous MDA5 expression. Expression of ISGs is compared between WT and ADAR1^E912A^ cells complemented with different ADARs. TPM (Transcripts Per Kilobase Million) is log2 transformed for the comparison. **d**, ISG induction in ADAR isoform-specific mutants. P150-specific exon is deleted in the p150 knockout. The p110 promoter is deleted to silence the p110 isoform (detail in methods). MDA5 is overexpressed in both mutant cell lines. **e**, Sub-cellular localization of ADAR proteins. ADAR proteins were tagged with Flag at the N-termini and immuno-stained with Flag antibody (left). DAPI staining is used to visualize the nucleus (middle). ADAR2 NLS sequence was removed to target the enzyme to the cytoplasm. **f**, ISG expression comparison between cytoplasmic ADAR2 complemented ADAR1 mutant cells and WT cells. **g**, Schematic diagram of the ADAR1–dsRNA–MDA5 axis showing cytosolic dsRNA editors (ADAR1p150/ADAR2△^NLS^) are required to suppress MDA5-mediated dsRNA sensing.

### ADAR1p150, but not p110, can efficiently suppress dsRNA immunity

Two protein isoforms are encoded by the *ADAR* gene; a long p150 isoform is predominantly cytoplasmic while a short p110 isoform is in the nucleus^25,27,29^. ADAR1p150 appears to be the key isoform suppressing the dsRNA-mediated immune response, since embryonic lethality of *Adar1p150*^*-/-*^ mice, which phenocopy *Adar1*^*-/-*^ mice^3^, can be rescued by deletion of *Mavs*^33^. To determine the role of ADAR1p150 in human cells, we lentivirally introduced ADAR1p150, ADAR1p110, and ADAR2 into the ADAR1^E912A^ cells with MDA5 expressed. ISG induction in the ADAR1^E912A^ cells was completely repressed by ADAR1p150, but not by ADAR1p110 or ADAR2, despite the fact that exogenous ADAR1p150 was expressed at lower levels than exogenous ADAR1p110 or ADAR2 (**Fig. 1c, Extended Data Fig. 2a**). Similarly, ADAR1p150 suppressed ISG induction in ADAR1^KO^ cells (**Extended Data Fig. 2b**). This complementation by ADAR1p150 is dependent on the RNA editing activity, as the deaminase inactive p150 mutant could not suppress the ISG induction (**Extended Data Fig. 2c**). In addition, we observed silencing of constitutively expressed MDA5 over time, so an inducible MDA5 was used for all subsequent experiments (**Extended Data Fig. 2d**).

To further confirm the functional difference between p150 and p110, we used HEK293T cells with loss of p150 (p150^KO^) or deficiency of p110 (p110^KD^) alone^40^. Upon MDA5 expression, we observed ISG induction in p150^KO^, but not in p110^KD^, cells (**Fig. 1d**). Taken together, our data strongly indicate that ADAR1p150 is the major isoform that suppresses the ISG induction in human cells, consistent with the *in vivo* mouse data.

### Cytoplasmic editing is required to suppress ISG induction

It is unclear why p150, but not p110, efficiently suppresses ISG induction. Both isoforms contain RNA binding and catalytic domains, but p150 harbors a unique Zα domain at the N terminus that binds left-handed Z-DNA/Z-RNA. Proteins with Zα domain have been implicated in immune-related functions^42,43^, which may explain p150’s unique function to prevent ISG induction. To test this, we generated a suite of p150 Zα mutants, including W195A, which is known to destabilize Zα^44^, and two additional mutants, P193A and G183S, which are observed in AGS patients^35,45^. In addition, we also included a deaminase domain mutant G1007R present in AGS patients^35^, and a deaminase deletion mutant as a positive control. All constructs were individually integrated into ADAR1 editing-deficient cells engineered to contain a Doxycycline-inducible MDA5 (**Extended Data Fig. 2d**). While mutations of the deaminase, particularly loss of the entire domain, impaired p150’s function in suppressing ISG induction, the mutations in Zα had no obvious effect (**Extended Data Fig. 2e**). Our finding appears to contradict with findings that suggest the importance of Zα domain of ADAR1 in human patients^35,46^ and animal models^47-54^. The discrepancy may suggest that overexpression of p150 in our cellular assays compensate the effect of the Zα domain.

An alternative explanation is the distinct cellular localization of each isoform. When expressed in HEK293T cells as FLAG-tagged proteins, ADAR1p110 and ADAR2 were predominantly in the nucleus while ADAR1p150 was mainly in the cytoplasm as indicated by the fluorescent immunostaining (**Fig. 1e**). To further distinguish the contributions of the Zα domain and localization for ADAR1 function, we deleted the nuclear localization signal (NLS) from ADAR2, which naturally lacks a Zα domain. This yielded an ADAR2△^NLS^ mutant that was mainly localized in the cytoplasm (**Fig. 1e**). Surprisingly, ADAR2△^NLS^ completely suppressed the ISG induction in ADAR1^E912A^ cells (**Fig. 1f**). Taken together, our results suggest that RNA editing in the cytoplasm is essential in suppressing MDA5-dependent dsRNA sensing (**Fig. 1g**).

### Identification of immunogenic dsRNAs by comparative RNA editing analysis

Based on our observation that cytoplasmic RNA editing is required to suppress ISG induction (**Fig. 1g**), we performed a comparative analysis of RNA editing by p150 vs p110 (or ADAR2 vs. ADAR2△^NLS^) to uncover the dsRNAs that are specifically, or preferentially, edited in the cytoplasm compared to the nucleus which we reasoned to be the key immunogenic dsRNA substrates (**Fig. 2a**).

**Fig. 2:**
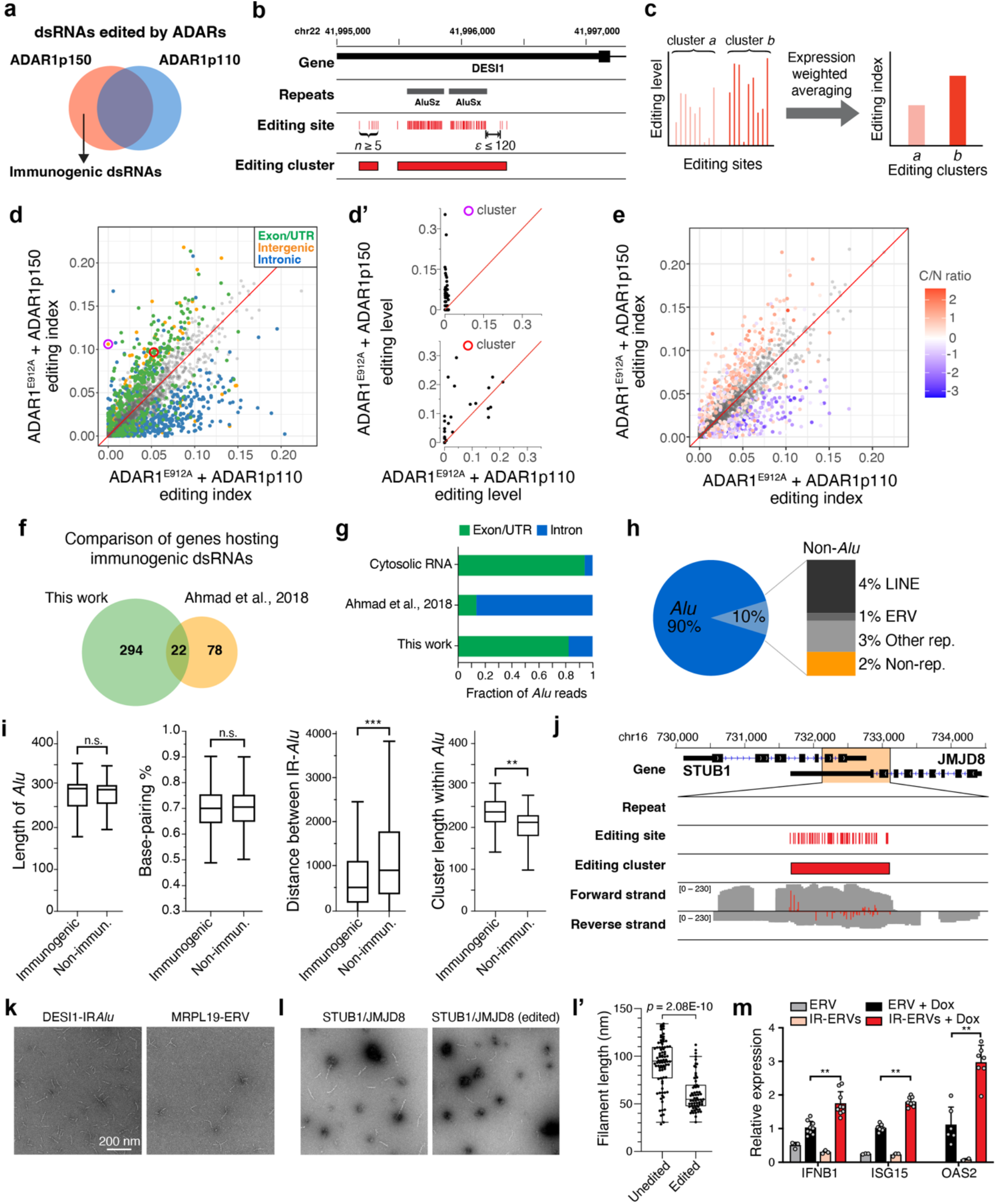
Identification and characterization of immunogenic dsRNAs in HEK293T cells. **a**, Schematic diagram showing identification of immunogenic dsRNAs through comparative analysis of ADAR substrates. **b**, Example of editing clusters. Editing sites within close proximity were classified as one cluster if (1) the number of editing sites (n) is at least 5 and (2) the distance between any two adjacent sites (ε) is less than 120 nt (see Methods for details). **c**, Schematic diagram of the conversion of editing levels at individual sites of a cluster to editing index of an entire editing cluster (see Methods for details). **d**, Editing index comparison between ADAR1^E912A^ cells complemented with ADAR1p110 and ADAR1p150. Clusters with significantly different editing index (adjusted p-value < 0.01) were colored based on genomic location (UTR, intergenic or intronic). **d’**, Editing level comparison of individual sites for two exemplary clusters indicated in purple and red circles in **d. e**, Editing clusters shown in **d** were color-coded according to the ratio of cytoplasmic and nuclear expression (C/N ratio) determined by RNA-seq of cytoplasmic and nuclear RNA (see Methods). **f**, Comparison of genes hosting Immunogenic dsRNAs identified in this study with the top 100 genes reported in a previous study^6^. **g**, Comparaive analysis of the RNA-seq reads mapped to *Alu*s in introns and exons/UTRs. RNA-seq data of the RNase protein assay (2.0 ng/μL RNase condition) was obtained from Ahmad et al., 2018^6^. Cytosolic RNA-seq data was used as control. **h**, Classification of immunogenic dsRNAs identified in HEK293T cells. **i**, Comparative analysis between immunogenic and non-immunogenic *Alu*s in *Alu* length, base-pairing percentage, distance between inverted *Alu*s (i.e., the loop size), and the length of the editing cluster within *Alu*. The non-immunogenic *Alu*s were randomly chosen from editing clusters that were expressed in exonic regions. **: *p* < 0.01, ***: *p* < 0.001, Student’s *t*-test. **j**, Example of a *cis*-NAT predicted to be a top ranked immunogenic dsRNA. Genes, repeats, editing sites, editing clusters, and strand-specific RNA-seq reads were shown in the expanded locus that spans the *cis*-NAT indicated by the editing cluster. **k**, Negative stain electron microscopy image of MDA5 filaments along the predicted dsRNAs (inverted repeat of two *Alu*s in the 3’UTR of *DESI1*, and two endogenous retroviruses (ERVs) in the 3’UTR of *MRPL19*). **l**, Negative stain electron microscopy image of MDA5 filaments along the *cis*-NAT formed by unedited (left) and edited (right) *STUB1* and *JMJD8* RNAs. **l’**, Comparison of the filament length between the unedited and edited *cis*-NAT. **m**, Real-time PCR measurement of three ISGs (*IFNB1, ISG15*, and *OAS2*) in ADAR1^E912A^ cells that were transfected with plasmids expressing single ERV or IR-ERV derived from *MRPL19* and treated with or without Dox to induce MDA5. Biological replicates n=2, technical replicates n ≥3. **: p < 0.01, Student’s *t*-test.

We initially compared the editing level of known editing sites (documented in the RADAR database^23^) individually using the RNA-seq data from the MDA5-expressing ADAR1^E912A^ HEK293T cells complemented with p150 or p110 (**Extended Data Fig. 3a**). We found that most of the sites were similarly edited by p150 and p110, and overall, p150-expressing cells had even lower level of editing than p110-expressing cells, probably due to the higher expression level of p110 than p150. This analysis made it challenging to identify individual RNA editing sites that are more highly edited by p150.

Comparing individual site editing is problematic as a way to identify immunogenic dsRNAs because editing of individual sites cannot reflect the overall editing status of the entire dsRNA that is often long and hyper-edited. Therefore, we developed a new strategy to compare RNA editing at the whole dsRNA level. To identify dsRNAs in an unbiased fashion, rather than relying on annotations such as *Alu* repeats, we performed *de novo* identification of editing clusters as a proxy of long dsRNAs (**Fig. 2b**). To define an editing cluster, we required that (1) it has at least five editing sites and (2) any two adjacent sites are less than 120 nt apart (see Methods, **Extended Data Fig. 3b–c**). Using this novel approach, we identified 205,652 clusters in HEK293T cells, ∼97% of which were in *Alu* and other repeats as exemplified in **Fig. 2b**. The remaining ∼3% of clusters in non-repeat regions were previously unannotated as dsRNA candidates. Importantly, identifying editing clusters allowed us to group all sites in a dsRNA to quantify the overall editing level, defined as the cluster editing index (see Methods) (**Fig. 2c**).

We performed a comparative analysis to search for editing clusters whose editing index is significantly higher in p150-than p110-complemented cells (**Fig. 2d**). We found that most (∼96%) of the editing clusters were similarly edited by p150 and p110, suggesting that most of the dsRNAs have no or low immunogenicity. Of 39,079 testable editing clusters with sufficient coverage, we identified 420 (1.1%) clusters located in 316 genes as putative immunogenic dsRNAs (**Table S1**), most of which were in the untranslated regions (UTRs) or intergenic regions but not in introns, consistent with the expectation that immunogenic dsRNAs recognized by cytosolic MDA5 would be embedded in processed RNAs exported to the cytoplasm. Of the clusters preferably edited by p150, some can be edited by p110 but to a lesser extent, while others are specifically edited by p150 only (**Fig. 2d’**). In contrast, editing clusters more highly edited by p110 were largely present in introns, probably as a result of the overexpression of p110 in the nucleus. These intronic dsRNAs would not be able to be sensed by cytosolic MDA5 as they are removed during RNA splicing, thus rendering them non-immunogenic.

We made similar findings when we compared cluster editing indices between ADAR2- and ADAR2△^NLS^-complemented cells (**Extended Data Fig. 3d**). The two lists of immunogenic dsRNAs identified from p150 vs. p110 or ADAR2 vs. ADAR2△^NLS^ comparisons were highly reproducible, particularly for immunogenic dsRNAs identified with higher statistical significance (**Extended Data Fig. 3e**).

In addition, we compared the cluster editing indices between WT and *ADAR1p150*^*-/-*^ cells to identify immunogenic dsRNAs that are significantly less edited in *ADAR1p150*^*-/-*^ cells. As expected, the immunogenic dsRNA candidates were enriched in the UTR and intergenic regions (**Extended Data Fig. 3f**). Furthermore, we performed RNA-seq on the cytoplasmic and nuclear RNA fractionations of wild type HEK293T cells (**Extended Data Fig. 3g**), and calculated the relative cytoplasmic-over-nuclear localization preference for each editing cluster (see Methods). We found that p150- and p110-preferred clusters were predominantly enriched in the cytoplasm and nucleus, respectively (**Fig. 2e**). These data confirm our findings that immunogenic dsRNAs are predominantly edited in the cytoplasm.

Finally, we compared our list of immunogenic dsRNAs identified in HEK293T cells with the recent finding from an *in vitro* biochemical assay in which dsRNAs purified from HEK293T cells were identified if they were bound by MDA5 filament and protected from RNase digestion^6^. We used all 316 immunogenic dsRNA-containing genes from our work (**Table S1**) and the top 100 dsRNA-containing genes annotated from the RNase protection assay. While *Alu* repeats were identified by both studies as the major source of immunogenicity, there were only 22 overlapping genes (**Fig. 2f**). We had p150 vs. p110 comparative RNA editing data for 73 of the 78 other genes in Ahmad et al.’s list; none of them met our statistical thresholds to be identified as immunogenic dsRNAs. Furthermore, our analysis of data from the RNase protection assay found that 82.6–91.5% of *Alu*-originated reads were mapped to intronic regions (**Fig. 2g, Extended Data Fig. 4a**), despite the fact that cytosolic RNA is used in the RNase protection assay^6^. Our analysis suggests that, in general, capture of immunogenic dsRNAs biochemically *in vitro* does not recapitulate the cellular conditions, leading to overly sensitive but non-specific binding of MDA5 to dsRNAs. Therefore, we instead examined if immunogenic dsRNA identified in this study were enriched in the RNase protection assay. We quantified the enrichment by the dsRNA expression fold-change in RNase-treated samples over untreated samples (in ADAR1 KO). We found that compared to control dsRNAs (randomly selected non-immunogenic dsRNAs located in 3’ UTR with comparable expression level to the immunogenic dsRNAs), the immunogenic dsRNAs were significantly more enriched for MDA5 binding (**Extended Data Fig. 4b**; median fold-change: 16.6±6.3 vs. 2.34±2.08, *p*-value = 4.065e-08, Wilcoxon signed-rank test).

### Characterization of immunogenic dsRNAs identified in HEK293T cells

Among the potentially immunogenic dsRNAs we identified, the vast majority were inverted repeats from the *Alu* family (90%) and other types of repetitive elements (8%) (**Fig. 2h**). We sought to characterize the sequence and structural features of immunogenic dsRNAs, with a focus on the most abundant *Alu* repeats. First, we found that although the intronic *Alu*s comprise the vast majority of all *Alu*s, they are almost excluded from the immunogenic group (**Fig. 2g**), which aligns with our recent independent work showing that over 90% of disease-relevant dsRNAs are located in exons and UTRs^13^. The enrichment of immunogenic dsRNAs in mature mRNAs is fully expected because they would be sensed by MDA5 in the cytoplasm. Our finding is in sharp contrast to studies utilizing *in vitro* RNase protection assay that yield mostly intronic IR*Alu*s as MDA5 substrates^6,7^. Second, we quantitatively compared the contribution of different dsRNA features for immunogenic vs. non-immunogenic *Alu* repeats (**Extended Data Fig. 5a**). We found no significant difference in the length of *Alu*s, the base-pairing percentage, or the expression level between the two groups (**Fig. 2i**). Instead, a feature that stood out was the distance between the arms of an inverted *Alu* pair (i.e., the loop size) (**Fig. 2i, Extended Data Fig. 5a**). Immunogenic inverted *Alu*s had significantly smaller loop sizes than their non-immunogenic counterparts. One possible explanation is that the secondary structure of a relatively long RNA is dynamic with optimal and suboptimal formations, and the inverted *Alu* dsRNA structure is more frequently favored when two *Alu*s are closer. To test this hypothesis, we compared the RNA structure ensemble and the frequency of forming the optimal inverted *Alu* dsRNA between the immunogenic and non-immunogenic *Alu*s. Indeed, the *Alu* pairs with longer loops tended to have more dynamic structural conformations, while the *Alu* pairs with shorter loops were more restricted to form the optimal dsRNA conformation with the minimal free energy (**Extended Data Fig. 5b**). This probably renders a dsRNA with two arms in closer proximity more stable and immunogenic upon MDA5 binding. In agreement with this, the immunogenic inverted *Alu*s tend to have longer editing cluster than the non-immunogenic ones (**Fig. 2i**). Our analysis suggests that the length and identity of *Alu*s, which are generally very similar to each other, contribute little to the immunogenicity of dsRNAs, but the distance between two *Alu* arms, which is highly variable, dictates the immunogenicity.

We next ranked the immunogenic dsRNAs identified in HEK293T cells based on the statistical significance of cytoplasmic vs. nuclear editing differences (see Methods). Unexpectedly, the top two ranked dsRNAs were located in non-repetitive regions that comprise only 2% of all the identified immunogenic dsRNAs (**Fig. 2h, Table S1**). These top dsRNAs are extensively edited by p150 but barely edited by p110 (**Tables S1 and S2**). They are 650 and 1539 bp in length (as measured by editing clusters), substantially longer than the typical length of *Alu*s (∼280 bp). We failed to identify inverted sequences such as *Alu* near the respective clusters that can form a long dsRNA. Instead, we found evidence to support that the dsRNA was formed by partially overlapping transcripts derived from two adjacent genes transcribed in opposite directions (**Fig. 2j**). These complementary transcripts, termed *cis*-natural antisense transcripts (*cis*-NATs)^55^, are transcribed in opposite directions from the same genomic locus, thus forming a perfect dsRNA when unedited. This model of dsRNA formation by *cis*-NATs is further supported by the strand-specific RNA-seq data and a cluster of A-to-G editing sites in the overlapping region only (**Fig. 2j**). These data suggest that while the vast majority of immunogenic dsRNAs are formed by inverted repeats, the *cis*-NATs represent a unique class of perfect paired dsRNAs that are likely to be highly immunogenic.

To validate immunogenic dsRNAs as MDA5 substrates, we examined their ability to form MDA5 filaments *in vitro* using negative stain electron microscopy. In agreement with previous findings^56^, we observed a correlation between dsRNA length and MDA5 filament length (**Extended Data Fig. 5c**). We then chose to validate candidate dsRNAs, representing inverted *Alu*, endogenous retroviral (ERV) repeats as well as *cis*-NATs. In all cases, we observed filaments with varying lengths dependent on the length of the dsRNA (**Fig. 2k–l, Extended Data Fig. 5d–e**). For the *cis*-NAT formed by JMJD8 and STUB1 overlapping 3’UTRs, we compared the filament formation between the unedited dsRNA and the dsRNA after *in vitro* editing by ADAR1. The *in vitro* editing status was confirmed by Sanger sequencing of 32 individual clones (**Extended Data Fig. 6**). The edited substrate resulted in a significant reduction in the length of the MDA5 filament compared to the unedited form (43% reduction on average, **Fig. 2l–l’**). The edited dsRNA could also form MDA5 filaments probably because the *in vitro* experimental conditions do not recapitulate cellular conditions. Nevertheless, these data validate the ability of various types of immunogenic dsRNAs to form a substrate capable of supporting MDA5 filament formation *in vitro*.

To validate the cellular immunogenicity of identified dsRNAs, we first attempted to knockdown endogenous dsRNAs – either *cis*-NATs or IR*Alu*s – in ADAR1^E912A^ cells with Dox-inducible MDA5 and a luciferase IFN reporter (Methods). Despite high knockdown efficiency, we observed negligible difference between the knockdown and non-targeting controls in cellular immune response (**Extended Data Fig. 5f–g**). We reasoned that this result was expected due to the redundancy of the 420 putatively immunogenic dsRNAs located in 316 genes in HEK293T cells (**Table S1**), which when not edited by ADAR1, collectively contribute to the MDA5-dependent IFN response. Thus, it is unlikely that knocking down an individual or even mulitple dsRNAs will generate any measurable difference in cellular immune response.

Given the challenge of using genetic perturbation to validate the putatively immunogenic dsRNAs, we evaluated their ability to trigger MDA5-dependent IFN response by overexpressing them, one at a time, in human cells. We selected two dsRNAs: one formed by inverted repeat ERVs (IR-ERVs) from the 3’UTR of *MRPL19* (**Extended Data Fig. 5h**) and the other by *STUB1:JMJD8 cis*-NATs (**Fig. 2j**). Particularly, to mimic the formation of *cis*-NATs, *STUB1* and *JMJD8* sequences were placed in the 3’UTR of mClover3 and mRuby3, respectively, that are co-expressed in a plasmid but in opposite directions. To control the immunogenicity induced by delivery methods, we included plasmids that expressed only ssRNA. Plasmids were introduced into the ADAR1^E912A^ cells with Dox-inducible MDA5. We hypothesized that, upon Dox treatment to induce MDA5, the ADAR1^E912A^ cells would elicit an even stronger IFN response if the overexpressed RNA is immunogenic. Indeed, we observed a significantly stronger MDA5-dependent IFN response, as indicated by the elevated expression of three signature ISGs, for both IR-ERVs dsRNAs (**Fig. 2m, Extended Data Fig. 5i**) and the *cis*-NAT (**Extended Data Fig. 5j–k**) compared to their respective ssRNA controls, thus validating the cellular immunogenicity of two candidate dsRNAs.

### Identification of immunogenic dsRNAs in human neural progenitor cells

Several lines of evidence suggest that there are different repertoires of immunogenic dsRNAs in different cell types, tissues, and organisms. While the *Adar1*^*-/-*^ and *Adar1*^*E861A/E861A*^ mice die of fetal liver/erythropoiesis failure, human AGS patients carrying ADAR1 loss-of-function mutations have severe brain inflammation^3,4,35,36^. The primate-specific *Alu* repeats, which consist of the vast majority of the edited dsRNAs in humans, are not conserved in mice. In addition, when *ADAR1*^*-/-*^ human embryonic stem cells (hESCs) were differentiated into neural progenitor cells (NPCs) or hepatocyte-like cells (HLCs), ISG induction and cell death phenotypes are only observed in NPCs, but not in HLCs^40^, consistent with brain inflammation being the major manifestation in human AGS patients.

To help uncover the potential immunogenic dsRNAs that are relevant in the human brain, we chose to use NPCs as a proof of concept. We differentiated the hESCs (both WT and *ADAR1*^*-/-*^)^40^ into NPCs (**Extended Data Fig. 7a–b**). Deletion of ADAR1 resulted in a severe cell death phenotype accompanied by the massive induction of ISGs at Day 29 after differentiation (**Fig. 3a, Extended Data Fig. 7c–d**). The ISG induction was not observed at Day 15 after differentiation (**Fig. 3a, Extended Data Fig. 7e**), although NPC marker PAX6 was expressed by this time point (**Extended Data Fig. 7b**).

**Fig. 3:**
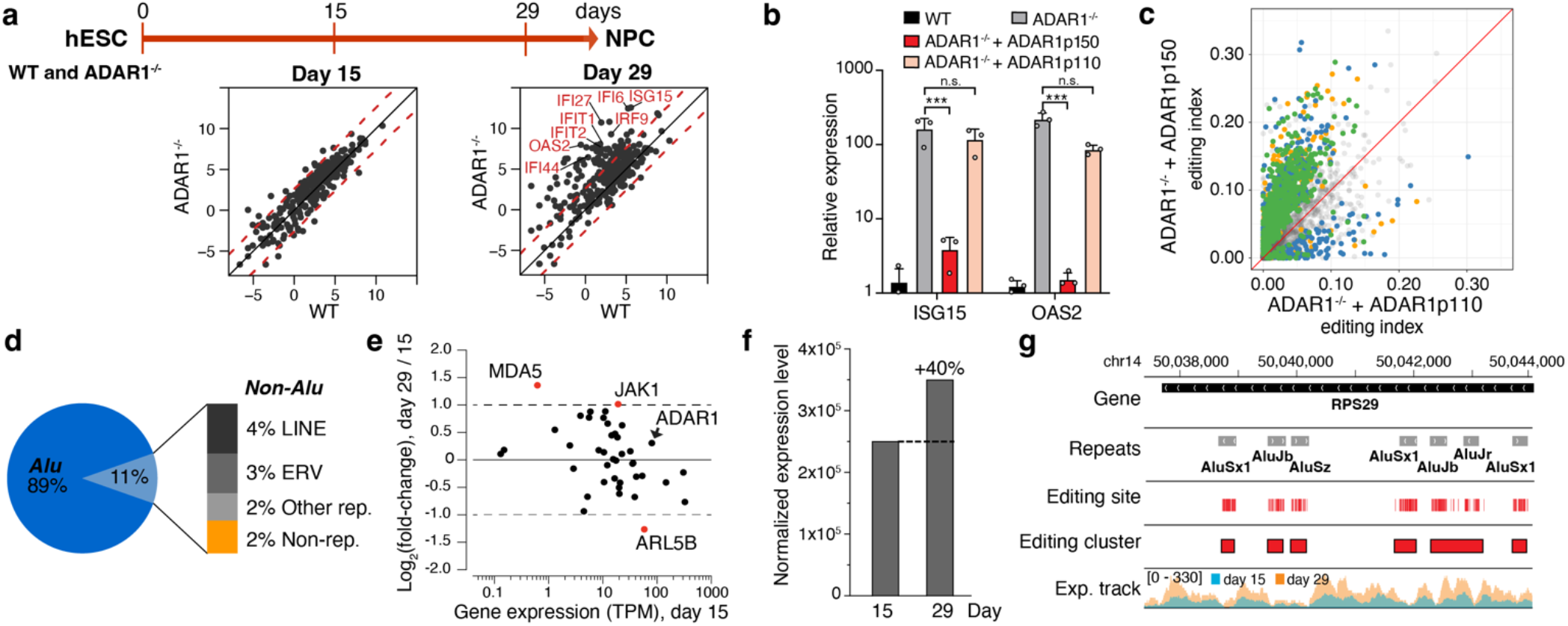
Identification and characterization of immunogenic dsRNAs in neural progenitor cells (NPCs). **a**, Comparison of ISG expression between WT and ADAR1^-/-^ NPCs at Day 15 and Day 29 after differentiation. Red dash lines indicate five-fold expression change. **b**, Signature ISG expression in hESC and NPC at Day 29 after differentiation. Real-time PCR was performed on RNA of WT and KO (ADAR1^-/-^) cells. Biological replicates n=3. n.s.: *p* ≥ 0.05, **: *p* < 0.01, Student’s *t*-test. **c**, Editing index comparison between ADAR1p110- and ADAR1p150-expressing ADAR1^-/-^ NPCs. Editing clusters with significantly different editing index (adjusted p-value < 0.01) were colored the same way as in **Fig. 2d. d**, Classification of immunogenic dsRNAs identified in NPCs. **e**, Expression change of 42 genes in the MDA5 and type-I IFN pathways between Day 15 and Day 29 post-differentiation of NPCs. **f**, The comparison of total expression levels of immunogenic dsRNAs (measured in TPM) between Day 15 and Day 29 post-differentiation of NPCs. **h**, Examples of immunogenic dsRNA editing status and expression levels between Day 15 and Day 29 post-differentiation of NPCs for *Alu*s. More examples for ERVs and *cis*-NATs are shown in **Extended Data Fig. 8**. Data were represented as mean ± SD for **b**.

To identify the potential immunogenic dsRNAs in the NPCs at Day 29 after differentiation from hESCs, we first performed RNA editing analysis using RNA-seq data from WT and *ADAR1*^*-/-*^ NPCs. Loss of ADAR1 led to a significant reduction of RNA editing in a large number of editing clusters (**Extended Data Fig. 7f**), although a substantial amount of editing remained due to the ADAR2 expression in NPCs. ADAR2-mediated editing in NPCs was further confirmed by the high level of editing of a *GRIA2* site, which is known to be specifically edited by ADAR2 (**Extended Data Fig. 7g)**. The presence of ADAR2, however, did not functionally compensate for the lack of ADAR1 to suppress the innate immune responses in NPCs, similar to our observations in HEK293T cells and previous *in vivo* mouse studies^57^.

To identify immunogenic dsRNAs in NPCs, we performed a comparative editing analysis similar to what we described for HEK293T cells. We stably integrated the p150 or p110 isoform into *ADAR1*^*-/-*^ NPCs immediately after the differentiation (**Extended Data Fig. 7h**). As expected, p150, but not p110, suppressed ISG induction (**Fig. 3b**). Of 40,182 testable editing clusters with sufficient coverage in NPCs, we identified 950 (2.4%) immunogenic dsRNA candidates that were significantly more highly edited by p150 than p110 (**Fig. 3c, Table S3**).

### Characterization of the immunogenic dsRNAs identified in NPCs

The categories of immunogenic dsRNAs identified in NPCs were similarly distributed to those identified in HEK293T cells, with 89% of them as *Alu* repeats (**Fig. 3d**). Again, we identified non-repetitive *cis*-NATs as the top ranked immunogenic dsRNAs candidates (**Table S2**). Compared to the editing clusters formed by *Alu*s, the editing clusters formed by LINE, LTR, and non-repetitive sequences were substantially longer, indicative of longer dsRNAs with potentially higher immunogenicity (**Extended Data Fig. 8a, Table S3**).

We next characterized features of immunogenic dsRNAs, again by focusing on the most abundant *Alu* repeats. We used a principal component analysis (PCA) approach to assess the importance of each dsRNA feature’s contribution to the variance of immunogenicity (see Methods for details). The immunogenic inverted *Alu*s from NPCs, like those in HEK293T cells, tended to have shorter distances between the two *Alu* arms, which may result in lower diversity of RNA structure ensemble and longer editing clusters within *Alu*s (**Extended Data Fig. 8b**).

The differentiated NPCs are a mixed population of neural cells. In a preliminary attempt to identify the relevant neural cell types in which the immunogenic dsRNAs are most likely to be expressed, we performed expression-weighted cell type enrichment analysis^58^ (see Methods). We found that NPC-specific immunogenic dsRNAs were most likely to be expressed in a selected group of neuron types and oligodendrocytes. In contrast, HEK293T-specific immunogenic dsRNAs showed no significant enrichment in any of the neural cell types (**Extended Data Fig. 8g**).

Finally, we investigated why the ISG induction and cell death of NPCs were observed at around Day 29, but not Day 15, after differentiation. First, we compared the expression of genes involved in interferon signaling pathways (**Table S4**) at Day 15 and Day 29 after differentiation in WT NPCs. We observed a 2.6-fold increase in MDA5 expression and 2.4-fold decrease in ARL5B expression at Day 29 compared to Day 15 (**Fig. 3e**). Interestingly, ARL5B is a known negative regulator of MDA5^59^. The other gene with greater than two-fold expression change was JAK1, which is essential for type I and II interferon signaling^60^. The expression changes of these interferon signaling proteins could potentially contribute to elevated ISG expression in NPCs at Day 29. Second, we examined the expression level of immunogenic dsRNAs identified in NPCs at Day 15 and Day 29. We found that, compared to Day 15, the total expression of immunogenic dsRNAs was 40% higher at Day 29 (**Fig. 3f**), as exemplified by *Alu*s (**Fig. 3g**), long intergenic LTRs (**Extended Data Fig. 8h**) and *cis*-NATs (**Extended Data Fig. 8i**). These data suggest that during NPC differentiation, the accumulation of immunogenic dsRNAs and/or the increased expression of interferon signaling proteins such as MDA5 activate the MDA5-mediated dsRNA sensing pathway and thus breach immune tolerance, implying a dose-dependent model.

### The cell type specificity of immunogenic dsRNAs

The immunogenic dsRNA candidates identified from HEK293T and NPC cells allowed us to understand their cell type specificity. Compared to the HEK293T cells, we identified two times more immunogenic dsRNAs in NPCs, suggesting the higher complexity of dsRNA editing and sensing in brain-related cell types (**Fig. 4a**). Only a subset (165) of immunogenic dsRNAs were found in both HEK293T and NPCs. When we confined our analysis to genes that are expressed in both cell types, the immunogenic dsRNA sets almost entirely overlapped (**Fig. 4a**). This suggests that the immunogenicity of a dsRNA is intrinsically determined by its sequence and structure features, and its effect is dependent on the host gene expression that may vary across cell and tissue types. This implies that the immunogenic dsRNAs identified in any cell types would be likely to be immunogenic in other cell types as long as they are expressed.

**Fig. 4:**
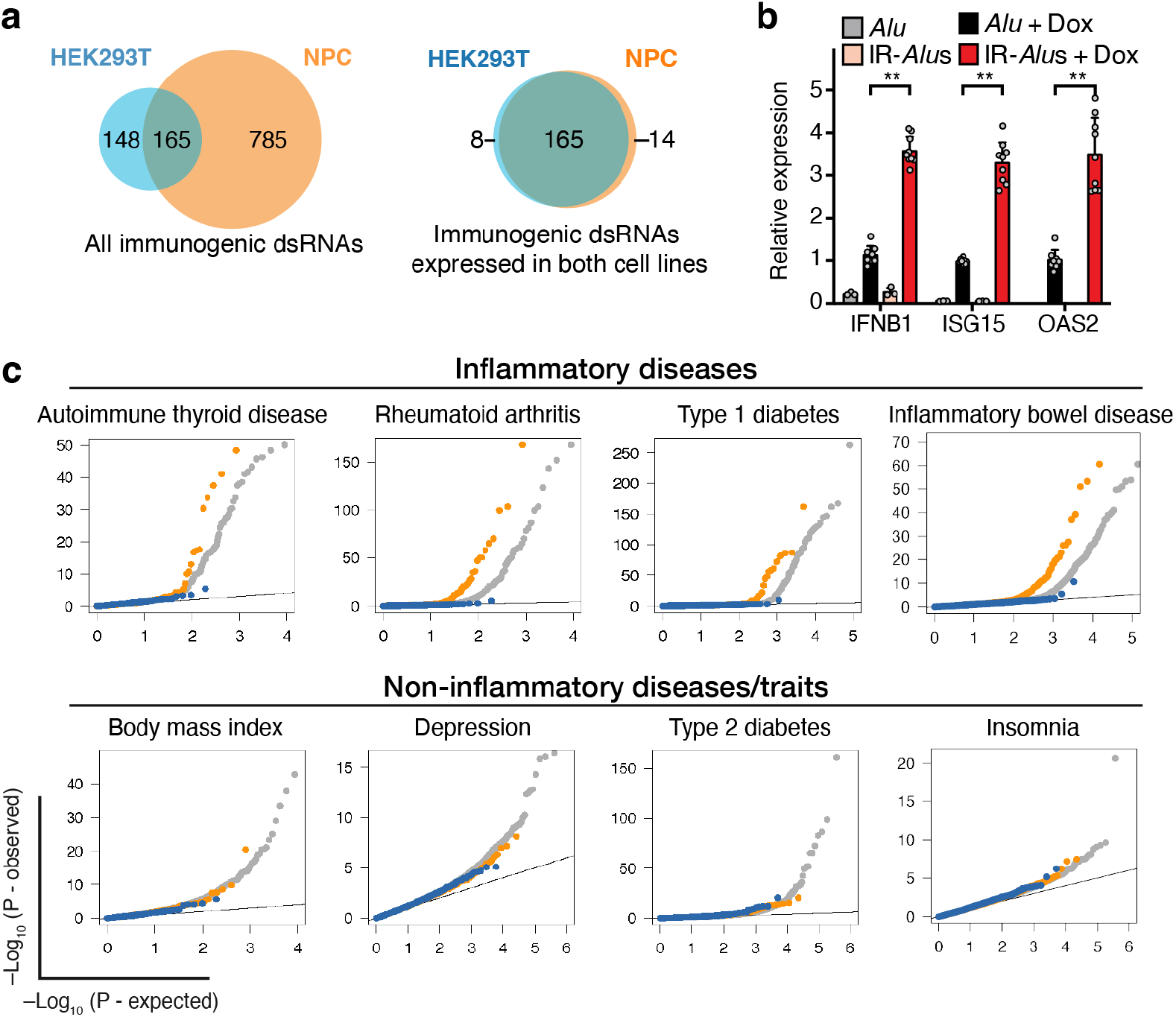
Immunogenic dsRNAs are cell type specific and disease relevant. **a**, Comparison of immunogenic dsRNAs identified in HEK293T and NPC cells (left) and those expressed in both cell lines (right). **b**, Real-time PCR measurement of the expression of three ISGs (IFNB1, ISG15, and OAS2) in ADAR1 editing-deficient HEK293T cells that were transfected with plasmids expressing single Alu or IR-Alus derived from NICN1 and treated with or without doxycycline (Dox) to induce MDA5. Biological replicates n=2, technical replicates n ≥3. **: p < 0.01, Student’s t-test. **c**, Quantile-Quantile (Q-Q) plots comparing enrichment for immunogenic dsRNAs (orange dots) vs. non-immunogenic dsRNAs (blue dots) in GWAS of 8 exemplary complex diseases and traits (grey dots). Top row: inflammatory diseases; bottom row: non-inflammatory diseases and traits.

We thus evaluated if an immunogenic dsRNAs identified in NPC only would be immunogenic when being expressed in HEK293T cells. We first chose an NPC-specific immunogenic dsRNA formed by IR-*Alu*s and overexpressed it in HEK293T cells, similar to the experiment described in **Fig. 2m** (see Methods). This IR-*Alu*s, compared to a single *Alu* as a negative control, led to an elevated IFN response that is dependent on MDA5 (**Fig. 4b, Extended Data Fig. 8c**). A similar effect was observed for an NPC-specific immunogenic dsRNA formed by *TAGLN:PCSK7 cis*-NAT when it was overexpressed in HEK293T cells (**Extended Data Fig. 8d–e**). To further test how the sequence and structure features may affect the immunogenicity of *cis*-NAT dsRNAs, we constructed a scrambled version of the *TAGLN:PCSK7 cis*-NAT that maintains the complementary structures of the overlapping region with randomized sequences (**Extended Data Fig. 8f**). We overexpressed this scrambled *cis*-NAT in the aforementioned HEK293T cells and observed elevated IFN response at comparable level to the original *cis*-NAT (**Extended Data Fig. 8d–e**). Our data suggests that the dsRNA structure, rather than its sequence, dictates the immunogenicity.

### Immunogenic dsRNAs are enriched at the GWAS signals of common inflammatory diseases

To genetically validate the immunogenicity of individual immunogenic dsRNA candidates, we attempted to silence the expression of a single dsRNA but failed to detect dampened immune responses (**Extended Data Fig. 5f–g**). This suggests that the contribution from an individual dsRNA is below the threshold to be measured by the conventional readout. Therefore, we took a different genetic approach to reveal the potential immunogenicity of the dsRNAs we identified, by taking advantage of the power of GWAS that can be very sensitive to detect weak signals with small effect sizes by using large sample sizes. We asked if immunogenic dsRNA candidates we identified would be enriched at the GWAS signals of common autoimmune and immune-related diseases, as we recently demonstrated in an independent project^13^. Indeed, we found that the immunogenic, but not the non-immunogenic dsRNA candidates, showed strong enrichment at the GWAS signals of inflammatory diseases, as exemplified in autoimmune thyroid disease, rheumatoid arthritis, type 1 diabetes, inflammatory bowel disease and others (**Fig. 4c, top row**; **Extended Data Fig. 9a**). Such enrichment, in contrast, was not observed for diseases and traits without apparent immune function implications, as exemplified by body mass index, depression, type 2 diabetes, insomnia, and others (**Fig. 4c, bottom row; Extended Data Fig. 9b**).

Our data suggests that immunogenic dsRNA candidates identified in this work, despite using HEK293T and NPC cells only, are potentially truly immunogenic and functionally relevant in inflammatory diseases. The experimental approach we developed in this work can be readily applied to disease-relevant cell types, which promises to uncover immunogenic dsRNAs that would be enriched at the GWAS signals of inflammatory diseases.

### The conservation of ADAR1-dsRNA-MDA5 axis between human and mouse

The ADAR1-dsRNA-MDA5 axis has been established with supporting evidence from mouse and human genetics. At the protein sequence level, ADAR1 and MDA5 are 76% and 80% identical between two species, respectively. However, it has not previously been determined whether ADAR1 and MDA5 are functionally interchangeable between human and mouse. Sequence conserved proteins are not necessarily functionally conserved across species.

To determine whether ADAR1 or MDA5 has acquired species specificity during evolution, we tested whether the mouse ADAR1 and MDA5 can substitute for their human counterparts in human cells. To test ADAR1, we transfected human or mouse ADAR1p150 into ADAR1^E912A^ HEK293T cells with Dox-inducible MDA5 and an IFN-responsive reporter (5xISRE-AP-NFkB-Lucia) to monitor the IFN induction (**Extended Data Fig. 10a–b**). Before the introduction of ADAR1p150, we observed MDA5-dependent activity of the IFN-responsive reporter (**Fig. 5a**). Upon transfection of either human or mouse ADAR1p150, the activity of the reporter was suppressed (**Fig. 5a**), suggesting that mouse ADAR1p150 can recognize and edit human immunogenic dsRNAs. To test MDA5, we stably integrated a human or mouse MDA5 in the human WT and *ADAR1*^*-/-*^ HEK293T cells (**Extended Data Fig. 10c**) and measured the expression of three signature ISGs. Both mouse and human MDA5 led to induction of ISGs in *ADAR1*^*-/-*^ cells (**Fig. 5b**). Taken together, our data provides evidence of functional interchangeability of ADAR1 and MDA5 between human and mouse.

**Fig. 5:**
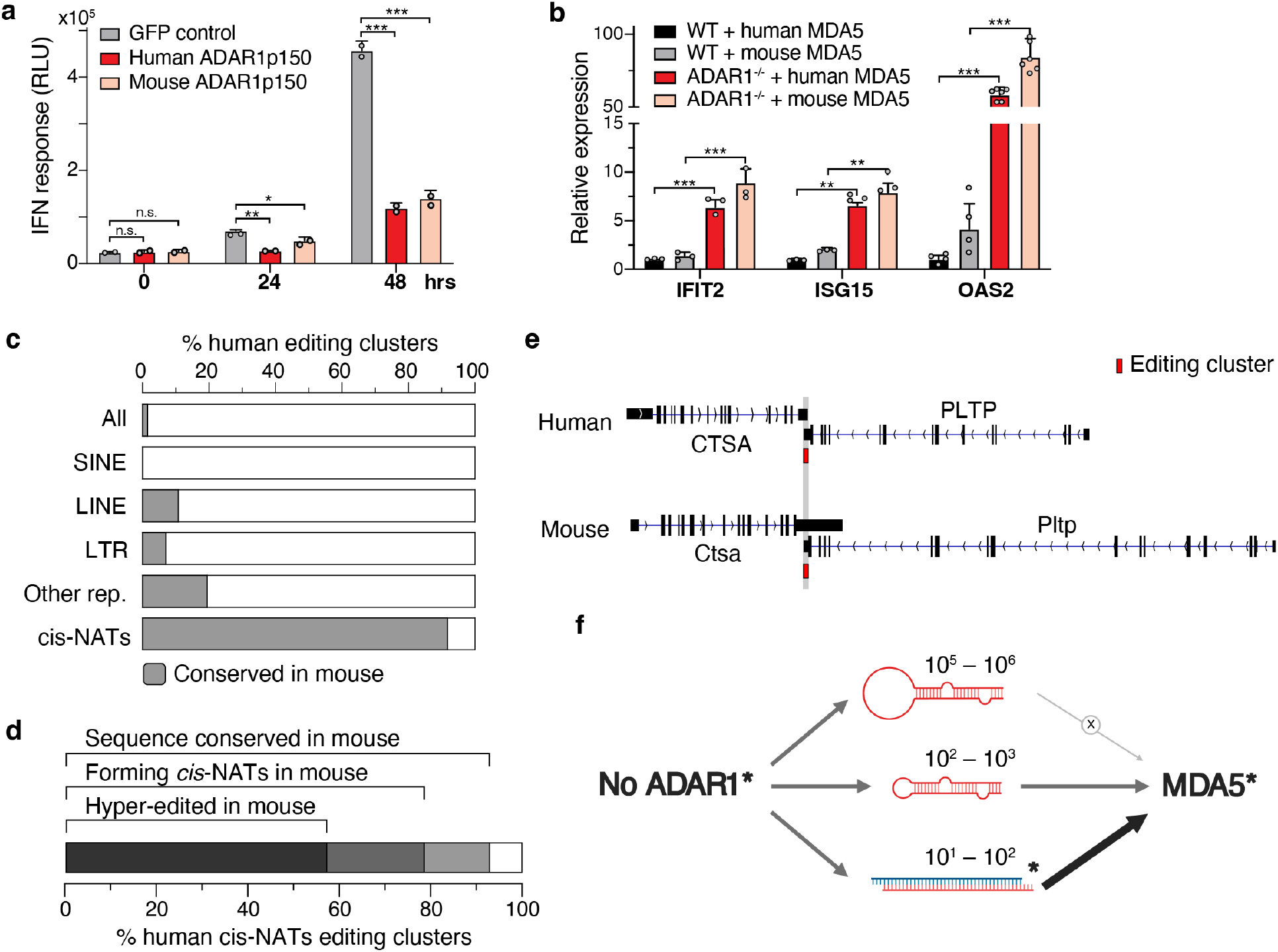
The conservation of the ADAR1-dsRNA-MDA5 axis between human and mouse. **a**, IFN response in ADAR1^E912A^ HEK293T cells transfected with human or mouse ADAR1p150. ADAR1^E912A^ cells with Dox-inducible MDA5 were treated with Dox for 0, 24, and 48 hours. IFN response was measured by luciferase activity in RLU (relative light unit) via a stably integrated ISRE-AP-NFkB-Lucia reporter in HEK293T cells (see Methods). GFP was used as a negative control. Biological replicates n=2. n.s.: *p* ≥ 0.05, *: *p* < 0.05, **: *p* < 0.01, ***: *p* < 0.001, Student’s *t*-test. **b**, Real-time PCR measurement of three ISGs (IFIT2, ISG15 and OAS2) in WT and ADAR1^-/-^ HEK293T cells that were stably integrated with human or mouse MDA5. Technical replicates n ≥3. **: *p* < 0.01, ***: *p* < 0.001, Student’s *t*-test. **c**, Fraction of human editing clusters of different classes that are conserved in the mouse genome. **d**, Fraction of immunogenic human *cis*-NATs that have supporting evidence to be conserved, also form *cis*-NATs and hyper-edited in mouse. **e**, An example of a conserved immunogenic *cis*-NAT between human and mouse formed by overlapping *CTSA* and *PLTP* genes. The editing cluster is indicated by the red box. **f**, Conservation of the ADAR1-dsRNA-MDA5 axis between human and mouse. Conserved components are labeled with *. Different groups of dsRNAs (with estimated total numbers shown), if not edited by ADAR1, have different levels of immunogenicity to activate MDA5. Data are represented as mean ± SD for **a** and **b**.

While ADAR1 and MDA5 are functionally conserved between human and mouse, dsRNA editing substrates have been regarded overall to be not^61-63^. When we examined all of the human immunogenic dsRNAs identified in this work, the dsRNAs residing in repeats were generally not conserved in mice (**Fig. 5c**). This is best demonstrated by the primate-specific *Alu*, a SINE element, where the vast majority of the millions of human editing sites reside. In contrast, of the 14 *cis*-NATs we identified in NPCs, 13 (93%) have a conserved genomic sequence in mice (**Fig. 5c, Table S2**). We then used a mouse brain RNA-seq dataset to determine whether the same *cis*-NATs form in mice (see Methods). We found supporting evidence that of the 14 identified human *cis*-NATs, 11 (78.6%) also formed in mice, and 8 (57%) were hyper-edited in mice (**Fig. 5d–e**). These percentages are likely underestimated because only one mouse RNA-seq dataset was used. Our data suggests that, while the majority of the immunogenic dsRNAs are not conserved across species, the *cis*-NATs represent a novel class of dsRNAs that tend to be conserved. The *cis*-NATs, despite relatively rare, are likely to be highly immunogenic compared to the abundant inverted repeats such as *Alu* (**Fig. 5f**).

## DISCUSSION

Innate immunity is the first line of defense against pathogens. MDA5, a cytosolic pattern recognition receptor, recognizes long dsRNAs, which are often produced through viral replication. How does MDA5 avoid sensing endogenous “self” RNAs that can frequently form dsRNA? The answer lies in RNA editing catalyzed by ADAR1, as demonstrated by both mouse and human genetics. Without sufficient RNA editing by ADAR1, un-or under-edited endogenous “self” dsRNAs can be recognized by MDA5 as “non-self”, triggering an interferon response. The discovery of the ADAR1-dsRNA-MDA5 axis points to the long dsRNAs, instead of protein-coding events, as the key ADAR1 substrates based on the substrate preference and signaling mechanism of MDA5. Identification of these immunogenic dsRNAs is very important because their impaired editing underlies the risk of both rare and common autoimmune and inflammatory diseases^13,35^.

There are a large number of endogenous dsRNAs in humans as indicated by the millions of human RNA editing sites and our editing cluster analysis. Is editing of each dsRNA equally important to evade MDA5-mediated immune responses? Based on the observation that cytosolic, but not nuclear, RNA editing suppresses MDA5-dependent immune response, we performed a comparative transcriptomic analysis to identify dsRNAs that are specifically or preferably edited by cytosolic vs. nuclear ADAR isoforms. While previous work has attempted to identify editing sites specifically edited by ADAR1 p150 vs p110^64,65^, there are two major advances in our work. First, instead of comparing editing levels at individual sites, we developed a novel method to calculate an editing index of each whole dsRNA, defined in an unbiased fashion as a cluster of editing sites. This is important because it is the editing status of the long dsRNA, rather than individual sites, that would influence the MDA5-dependent immunogenicity. Second, we used MDA5-dependent IFN response as the functional readout. This allowed us to enrich immunogenic dsRNAs whose insufficient editing would result in immune responses. We found that cytosolic editing of a small subset of dsRNAs (estimated 1.1% in HEK293T cells and 2.4% in NPCs), which we call immunogenic dsRNAs, is particularly important to avoid interferon response. Despite the relatively small fraction, the total number of immunogenic dsRNAs is still in the order of hundreds, with each contributing a small proportion to the total cellular immune response mediated by MDA5. Future work is needed to provide direct evidence of the immunogenicity of individual dsRNA candidates, although it may be compromised by the low specificity of the biochemical approaches (see below). In addition, we cannot conclude that the immunogenic dsRNAs identified in this study constitute the complete repertoire of the investigated cell types. Additional endogenous dsRNAs may also contribute to MDA5 activation to a lesser extent.

The immunogenic dsRNAs are largely located in the UTRs, although the vast majority of endogenous dsRNAs are located in introns. This is in agreement with the fact that immunogenic dsRNAs must be cytoplasmic to be sensed by MDA5. In addition to inverted repeats such as the prevalent *Alu*s, we discovered and validated other types of immunogenic dsRNAs such as Inverted ERV (**Fig. 2m**). Another previously unidentified dsRNA type is *cis-*NATs that are formed by overlapping genes transcribed in opposite directions. *cis-*NATs are rare; there were 2 in HEK293T cells and 14 in NPCs, all of which were identified as immunogenic dsRNA. However, their potential immunogenicity can be very high because they can form long, perfect dsRNAs. It is noteworthy that *cis-*NATs are highly enriched near risk loci of many inflammatory diseases^13^, further suggesting their functional importance. Activation of MDA5 by intermolecular dsRNA has been reported; dsRNAs formed by bi-directionally transcribed endogenous retrovirus elements or mitochondrial heavy and light strands can trigger MDA5-mediated interferon response^66-69^. Similarly in cancer cells, certain 3’ UTR of coding genes and antisense ERVs could form intermolecular dsRNAs that activate downstream IFN immune response^70^. For inverted *Alu* repeats, the key feature that distinguishes the immunogenic from non-immunogenic dsRNAs is the distance, but not the sequence similarity, between the two repeats comprising the *Alu* pair (**Fig. 2i, Extended Data Fig. 5a–b, Extended Data Fig. 8b**). This is consistent with the previous finding that the editability of *Alu* is mostly affected by the distance, but not similarity, of two closest reversely oriented neighboring *Alu*s^71^. This is likely due to the fact that *Alu*s are generally highly similar, but the distance between the members of *Alu* pairs is variable. A longer distance possibly makes it more difficult for two *Alu*s to fold back with each other, making the inverted *Alu* dsRNA conformation less favorable. Further work is needed to uncover the mechanism by which some dsRNAs are more immunogenic and how these immunogenic dsRNAs evade nuclear editing and are preferentially edited in the cytoplasm.

Our genetic approach to identifying the immunogenic dsRNAs in a cellular context has advantages over the biochemical approach of identifying dsRNAs bound by MDA5 filament *in vitro*^6^. Our analysis indicates that the ability of MDA5 to distinguish cellular immunogenic from non-immunogenic dsRNAs is poorly recapitulated in a test tube. First, the vast majority of the dsRNAs identified by the biochemical approach are located in introns, thus unlikely to be substrates of cytosolic MDA5 (**Fig. 2g, Extended Data Fig. 4a**). Second, they are not differentially edited in the cytoplasm vs. nucleus (**Fig. 2f**). Third, immunogenic dsRNAs, even when fully edited, can form MDA5 filaments *in vitro* (**Fig. 2l**).

We applied our genetic approach to identifying immunogenic dsRNAs, first in HEK293T cells, and then in NPCs that are more physiologically relevant because of the brain inflammation seen in AGS patients with ADAR1 LOF or MDA5 GOF mutations^35,37^. We identified almost identical sets of immunogenic dsRNAs in HEK293T cells and NPCs when considering genes that are expressed in both cell types. When overexpressed in HEK293T cells, NPC-specific immunogenic dsRNAs (either IR*Alu* or *cis*-NAT) could also trigger an immune response, implying that intrinsic sequence and structure features dictate the immunogenicity of a dsRNA. Many more immunogenic dsRNAs were present in genes that are only expressed in NPCs in an accumulative fashion during differentiation enabling them to reach a threshold of dsRNAs triggering immune responses, suggesting a need for a spatiotemporal search for immunogenic dsRNAs. We expect that our approach will be applied to the identification of immunogenic dsRNAs in different human cell types, as a necessary step to reveal the probable dsRNAs involved in autoimmune and inflammatory diseases^13^.

In another recent work of ours, we uncovered hundreds of immunogenic dsRNA candidates that are colocalized at the GWAS loci for autoimmune and immune-related diseases^13^. While this computational approach is successful in identifying disease relevant dsRNAs, it is limited by the availability of natural variations near dsRNAs in the human population. Our current method, which does not rely on the natural variations, enables an unbiased search of immunogenic dsRNAs in a given cell type. It is noteworthy that the immunogenic dsRNA candidates identified in this work appear to contribute to the inflammatory diseases (**Fig. 4c, Extended Data Fig. 9**), which strongly suggests their immunogenicity. Our method, when applied to many disease-related cell types in the future, promises to move us towards a comprehensive map of immunogenic dsRNAs that are relevant in human inflammatory diseases.

In addition to cell-type specificity, immunogenic dsRNAs differ across species. The primate-specific *Alu* and rodent-specific B1/B2 SINE elements are independently inserted in the respective genomes, thus making it highly unlikely that they share the same host genes. In contrast to the SINE repeats, the *cis-* NATs revealed in our work are generally conserved between human and mouse (**Fig. 5c–e**). Furthermore, we confirmed the functional interchangeablity of ADAR1 and MDA5 between human and mouse. Therefore, the ADAR1-dsRNA-MDA5 axis is highly conserved in general, with a major distinction being the particular immunogenic dsRNAs formed by the abundant SINE elements.

There are therapeutic potentials to harnessing immunogenic dsRNAs for treating human diseases. On one hand, as the immunogenic dsRNAs trigger immune activation, silencing them at the transcriptional level could dampen the chronic innate immune activation in some types of autoimmune disease. On the other hand, activating the ADAR1-dsRNA-MDA5 axis by delivering or upregulating immunogenic dsRNAs in tumors could be beneficial for cancer patients. Some cancer cell lines are sensitive to loss of ADAR1 due to the accumulation of unedited dsRNAs^8-10^. Recent studies showed that epigenetic therapy, such as inhibition of DNA methylation using 5-Aza-CdR and ablation of histone demethylase LSD1, leads to dsRNA accumulation, which subsequently triggers MDA5-mediated immune responses and slows down cancer cell growth^66,67,69^. Particularly, ADAR1 depletion could further sensitizes tumors to 5-Aza-CdR treatment^7^. These approaches to activating dsRNA sensing, via epigenetic therapy and/or ADAR1 silencing, can overcome resistance to immune checkpoint blockade such as anti-CTLA4 or anti-PD1 in mouse models^72^, which is analogous to activation of the dsDNA-mediated cGAS/STING innate immune pathway^73^. The immunogenic dsRNAs identified in this work may be exploited as specific agonists of MDA5 to turn an immunologically “cold” tumor “hot”, thus making tumors sensitive to immunotherapy. Therefore, our work fills an important knowledge gap in the ADAR1-dsRNA-MDA5 axis, and sheds light on both mechanistic understanding and potential therapeutic interventions.

## Methods

### Cell Lines

HEK293T cells were maintained in DMEM (High glucose, L-glutamine, Pyruvate) (Thermo Fisher Scientific) supplemented with 10% fetal bovine serum, 1% penicillin/streptomycin. Human ESCs (wild type and ADAR1 KO, gifts from Dr. C. Rice, the Rockefeller University) were maintained in complete mTeSR1 (StemCell Technologies) and the differentiated neural progenitor cells were maintained in STEMdiff Neural Progenitor Medium (StemCell Technologies). All cells were grown at 37°C in a 5% CO_2_ humidified atmosphere. The sex of HEK293T and hESC cell lines is female and male, respectively.

### Construct generation

Cas9 nickase vector px335-SpCas9n-D10A (Addgene #42335) and px462-SpCas9n-2A-Puro-D10A (Addgene #62987) were used as backbones for ADAR1 knockin and knockout. The guide RNA pairs were predicted at crispr.mit.edu. The backbone vectors were linearized by BbsI restriction digestion and ligated to the annealed oligos following the previously published method. The guide RNA pair was subsequently cloned into the same backbone by amplifying one cassette using primers px335-1F-NcoI and px335-730R and ligating it into the NcoI site. MAVS knockout was generated using the Cas9 vector px330 (Addgene #42230).

Protein overexpression constructs were generated by amplification of CDS of ADAR1 or ADAR2 or MDA5 and integration into the pCDH-puro backbone. Mouse ADAR1 was cloned into the backbone of the pAAV-CAG-GFP vector. Cytoplasmic ADAR2 mutant was generated by deletion of the first 70 amino acids coding sequence that encompasses the NLS of the ADAR2 CDS. ADAR1 point mutations were generated based on the ADAR1p150 sequence using the QuickChange II XL site-directed mutagenesis kit (Agilent). The doxycycline-inducible MDA5 construct was generated by replacing dCas9 with MDA5 CDS in the HR-TRE3G-dCas9-GCN4-10x-p2a-mCherry backbone. For protein purification constructs, human MDA5 was cloned into pET-50b vector, which contains an N-terminal His6-NusA tag followed by a HRV3C cleavage site. RNA overexpression constructs were generated by amplification of the ERV or *Alu* repeats and integration into the 3’UTR of the pCDH-EGFP.

### Generation of CRISPR mutant cell lines

HEK293T cells were maintained in DMEM media supplemented with 10% FBS and penicillin/streptomycin. 7×10^5^ cells were seeded one day before transfection of CRISPR constructs. For the ADAR1 E912A mutation, 600 ng guide RNA pair and Cas9 nickase (with both guide RNAs in the px335 backbone) along with donor oligo ultramer-912 were transfected into the cell using lipofectamine 2000 following the manufacturer’s manual. Fresh media with 10 μM L755507 (SML1362, Sigma-Aldrich) were added to the cells 6 hours after the transfection. The cells were cultured for 5 days with regular splitting before single clone selection using the dilution method. Briefly, 1000 cells were seeded on the 15 cm^2^ dish and cultured until the colonies were visible. Single clones were picked and transferred to 24-well plates with DMEM/FBS media. A small fraction of each colony was harvested and resuspended in squishing buffer (10 mM Tris-HCl pH 8.2, 1 mM EDTA, 25 mM NaCl and 200 μg/ml Proteinase K). The cells were incubated at 37°C for 30 minutes and 94°C for 5 minutes before direct usage for genotype screen using an ADAR-912-KI (or WT)-allele-specific primer (**Table S5**). The genotype of positive clones was further confirmed by amplifying the genomic region spanning -250nt to +250nt and generation of TOPO clones using TOPO PCR2.1 kit. TOPO clones were subject to Sanger sequencing at Quintara Biosciences (San Francisco, CA). Heterozygous clones with both the E912A allele and wild type alleles were selected for the next round of knock-in. Three rounds of knock-in were performed to obtain the homozygous E912A mutant. ADAR1^KO^ and ADAR1^E912A^;MAVS^KO^ HEK293T mutants were generated and selected using a similar approach as the above. In addition, HEK293T ADAR1p150^KO^ and ADAR1p110^KD^ were obtained from C. Rice lab (The Rockefeller University)^40^.

### RNA extraction and reverse transcription

Approximately 2-5 million HEK293T wild type or mutant cells were collected and the RNA was extracted using the Zymo RNA miniprep kit following the manufacturer’s manual. Genomic DNA was removed by on-column digestion of DNase I. Human embryonic stem cells (hESCs) and the differentiated neural progenitor cells (NPCs) were collected and RNA was extracted using the Trizol method. Briefly, cells were resuspended in 500-1000 μl Trizol and incubated at room temperature for 5 minutes, and debris was removed after centrifugation. 0.2 volumes of chloroform were added to the supernatant. The aqueous phase was collected after centrifugation at 4°C and mixed with 1 volume of isopropanol at -20°C for at least 30 minutes. RNA was precipitated at 13,000 rpm at 4°C. 1-2 μg RNA was used for cDNA synthesis using the iScript Advanced kit (Bio-Rad), following the manual.

### Nuclear and cytosolic fractionation

For nuclear and cytosolic RNA purification, nuclear and cytosolic fractionation kit (Cell Biolabs: AKR-171) was used. Briefly, the cytosolic fraction was released using hypotonic buffer without disturbing the nucleus. Nucleus was washed and then extracted using nuclear extraction buffer. The nuclear and cytosolic fractions were further extracted for RNA using the Trizol method.

For nuclear and cytosolic proteins, cells underwent a double thymidine block using 2mM thymidine (Sigma) to synchronize cells in G1/S phase and minimize the occurrence of nuclear-cytoplasmic mixing. Cells with each ADAR isoform were fractionated with the Cell Fractionation Kit (Abcam: ab109719) according to the manufacturer’s instructions. 5% of the resulting lysate from the nuclear and cytoplasmic fractions were resolved by SDS-PAGE gel electrophoresis using 4-15% gradient gels (Bio-rad) and transferred to nitrocellulose membranes. Primary antibodies used were mouse anti-FLAG (Sigma: F3165; 1:5,000), mouse anti-GAPDH (Thermo Fisher: GA1R; 1:5,000), and rabbit polyclonal anti-Histone H1 (PA5-30055). Secondary antibody incubation was carried out using the following antibodies diluted in PBS-T: goat anti-mouse-HRP (1:10,000) and goat anti-rabbit-HRP (1:10,000). Following three washes with PBS-T, the blot was developed with Pierce ECL Plus Western Blotting Substrate (Thermo Fisher 32132). GAPDH and Histone H1 were probed on separate blots.

### Real-time PCR

cDNA synthesized using the iScript Advanced kit was diluted 3-5 fold with nuclease-free H_2_O. KapaFast qPCR mix (2x) was used for real-time PCR. Primer sequences for each tested gene were obtained from qPrimerDepot and Primer Bank (https://pga.mgh.harvard.edu/primerbank/). 200nM primers were used in each reaction and run on the Bio-Rad CFX96 with the user manual program.

### RNA-seq library preparation and sequencing

rRNA was depleted from total RNA using RNase H-based protocols^74,75^. In brief, approximately 100-1000 ng of RNA was annealed with pooled rRNA antisense DNA oligo (gift from J. Salzman lab at Stanford) by heating at 94°C and slowly cooling down before Hybridase Thermostable RNase H (Epicenter, Madison, WI) incubation at 45°C for 30 minutes. Excessive DNA oligo was removed by treatment with TURBO DNase (Invitrogen: AM1907). RNA purification was performed using Agencourt RNAClean XP beads (Beckman Coulter: A63987). RNAseq libraries were prepared using the KAPA HyperPrep RNA-seq Kit (Kapa Biosystems: KK8540). RNAseq libraries were sequenced on the NextSeq (Stanford) or HiSeq (Novogene, CA). Sequencing parameters are provided along with the deposited data.

### Inducible MDA5 cell line generation

Wild type HEK293T cells were maintained in DMEM supplemented with 10% FBS and penicillin/streptomycin. Lentivirus expressing rtTA and TRE-MDA5-P2A-mCherry were separately packaged using the 2^nd^ generation system in HEK293T cells. The two viruses were co-infected into wild type or ADAR1^E912A^ HEK293T cells with 8 μg/ml polybrene. 2 μg/ml doxycycline was added to the media >48 hours post-infection. mCherry-positive cells were FACS sorted (at Stanford FACS facility) and cultured in doxycycline-free media to turn off MDA5 expression.

### ADAR1 mutant cells complementation assay

ADAR1 KO, ADAR1^E912A^, or ADAR1^E912A^ with inducible MDA5 cells were maintained in DMEM with 10% and penicillin/streptomycin. Lentivirus expressing CMV promoter-driven ADAR1, ADAR2 or MDA5 was packaged in HEK293T cells. Cells were infected with low titer virus expressing ADAR genes and were selected by 2 μg/ml puromycin. Equal amount of MDA5-expressing virus was used to infect the above ADAR-overexpressing cells.

### Immunofluorescence staining

HEK293T cells with stably integrated CMV-Flag-ADAR1 or CMV-Flag-ADAR2 were used in this assay. To visualize localization of different Flag-tagged ADAR proteins, cells were fixed with 1% PFA for 15 minutes at room temperature, followed by permeabilization with 0.5% Triton X-100. Then, cells were stained with Flag antibody (Sigma) for 1 hour and then incubated with secondary antibody. Nuclei were counterstained with DAPI. Images were taken using Leica DM RXA2.

### MDA5 protein expression and purification

Recombinant pET50b plasmids containing cDNA of HsMDA5 (298-1025) were transformed into *E. coli* C41 cells, which were cultured in LB medium with an extra 0.2 mM ZnCl_2_ at 37°C until OD_600_ reached 0.6-0.8. Protein expression was induced by adding 0.2 mM IPTG. After 20 h incubation at 18°C with shaking, the cells were harvested by centrifugation, resuspended in a buffer containing 20 mM Tris-HCl, 500 mM NaCl, 5% glycerol, 25 mM imidazole, 0.5 mM PMSF, pH 8.0 on ice, and lysed with high-pressure homogenization. Proteins were purified to homogeneity using Ni-NTA affinity chromatography, cation exchange, a second Ni-NTA affinity chromatography to remove the His6-NusA tag, and size exclusion chromatography (in that order). HRV3C protease was added after the first Ni-NTA affinity chromatography for cleavage of His6-NusA ^76^.

### Preparation of dsRNA *in vitro*

All dsRNAs were in vitro transcribed using T7 RNA polymerase. The two complementary strands were transcribed and purified separately. pUC19 plasmids containing target sequences were linearized by EcoRI, extracted with phenol chloroform and precipitated with isopropanol. In vitro transcription reactions were carried out at 37°C for 4 h in buffer containing 100 mM HEPES-K (pH 7.9), 10 mM MgCl_2_, 10 mM DTT, 6mM each NTP, 2 mM Spermidine, 200 μg/mL linearized plasmid, and 100 μg/mL T7 RNA polymerase. DNA template was digested with DNase I. Transcripts were purified by 8% denaturing urea PAGE, eluted from gel slices and precipitated with isopropanol. Prior to filament formation, complementary strands were mixed at a molar ratio of 1:1 in annealing buffer (10 mM Tris-HCl pH 7.5, 50 mM NaCl, 1 mM EDTA) and heated to 90°C for 2 min, then slowly cooled to room temperature.

### Negative stain electron microscopy

Samples including 0.32 μM HsMDA5(298-1025) and 2 ng/μL dsRNA (regardless of length) were incubated at room temperature for 20 min with 1 mM ADP▪AlF4 in buffer containing 20 mM Tris-HCl, 100 mM NaCl, 2 mM MgCl_2_, 2 mM DTT, pH 8.0. 5 μl of each sample was applied to glow-discharged 300 mesh carbon-coated copper grids (Beijing Zhongjingkeyi Technology), stained with 0.75% uranyl formate and air-dried. Data were collected on a Talos L120C transmission electron microscope equipped with a 4K × 4K CETA CCD camera (FEI). Images were recorded at a nominal magnification of 57,000×, corresponding to a pixel size of 2.5 Å per pixel. Filament lengths were measured using ImageJ.

### Cas13d-based chemiluminescence assay for measuring IFN response

Cas13d guides targeting the 3’UTR of individual *cis*-NAT pair members were selected from https://cas13design.nygenome.org/ and cloned into a plasmid bearing a guide expression cassette under the mouse U6 promoter and an mCherry-P2A-RfxCas13d expression cassette under the control of the EF1a promoter (a kind gift from LS Qi). The RfxCas13d additionally bears two nuclear localization signals and three FLAG tags. 125 ng of each plasmid was transfected using Lipofectamine 3000 into HEK293T-ADAR1-E912A-iMDA5-mCherry-pIFN-Lucia cells plated 2 days prior in a 96-well plate at 15,000 cells per well. 24 hours post-transfection, media was aspirated and replaced with 100 uL of growth media containing either 0 or 0.1 ug/mL of doxycycline. 24 hours post-transfection, 20 uL of supernatant from each well was collected and mixed with 50 uL of Quanti-Luc Gold solution (rep-qlcg1, Invivogen) in a white 96-well microplate (655075, Greiner Bio-One) before imaging on a Varioskan Lux plate reader (ThermoScientific) under the following conditions: 1 second shaking, 1000ms exposure collection. For each plate, a minimum of 3 replicates per condition were used.

### dsRNA candidate overexpression assay

EGFP constructs with inverted repeat dsRNA candidates in the 3’UTR were packaged into lentivirus and used to infect HEK293T ADAR1^E912A^ cells with inducible MDA5. Equal amounts of virus were used for the putative inverted repeat dsRNA and its control. Two days after the infection, cells were treated with 1μg/ml Dox for another two days to induce MDA5. Cells were harvested for RT-qPCR analysis of signature ISGs. For the *STUB1:JMJD8* and *TAGLN:PCSK7 cis*-NATs as well as the scramble control, plasmids were constructed convergently expressing mClover3 and mRuby3 under the control of EF1a promoters with both genes’ 3’ termini consisting of the overlapping genomic sequence of the relevant *cis*-NAT pair (**Extended Data Fig. 8f**). These plasmids were transfected into HEK293T ADAR1^E912A^ cells with inducible MDA5 plated 24 hours prior at 125,000-150,000 cells per well in 24-well tissue culture plates. Transfection was carried out using 500 ng of plasmid and Lipofectamine 3000 in a 3:1 lipofectamine:plasmid amount ratio. 24 hours post-transfection, media was replaced with media containing 0.1μg/mL doxycycline. 24 hours post Dox addition, cells were harvested for RT-qPCR analysis of signature ISGs.

### Luciferase assay

ISRE-AP-NFkB-Lucia reporter was originally purchased from Invivogen and was stably integrated into HEK293T ADAR1^E912A^ cells. Inducible MDA5-P2A-mCherry was also stably expressed in the same cell. 1×10^6^ cells were seeded in each well of the 6-well plate one day before the transfection of the ADAR1 or GFP constructs. 500ng of EGFP or mouse ADAR1p150-EGFP or human ADAR1p150 were transfected into the cells using lipofectamine 2000 (n=2). 6 hours post transfection, the media was changed, and 2μg/ml doxycycline was added. 20μl of media was harvested at 0, 24, and 48 hours post transfection. Media was mixed with Gold-Lucia substrate (Invivogen) before measurement using a plate reader.

### NPC differentiation and complementation

Wild type and ADAR1^-/-^ human ESCs were grown in mTeSR1 media (STEMCELL) before differentiation into NPCs following the STEMCELL manual. In brief, cells were induced in NPC induction media (STEMCELL, 05835) with 10μM Y-27632 for 13 days and were split when they reached confluency. After induction, cells were maintained in neural progenitor media (STEMCELL, 05833) until ADAR1^-/-^ cells started to exhibit cell death phenotype. The bright-field images were captured by SI8000 Cell Motion Imaging System (Sony Biotechnology) using a 4x objective. For the genetic complementation assay, CMV-ADAR1p110 or CMV-ADAR1p150 was lentivirally integrated into ADAR1^-/-^ NPCs at day 17 post-induction and stable cells were selected by puromycin at day 19. Complemented NPCs were maintained regularly until day 29. Cells were harvested and RNA was extracted using Trizol and chloroform first, and then the aqueous phase was passed through the RNeasy column (Qiagen) for RNA purification.

### Mapping of RNA-seq reads

We adopted our previously published pipeline to accurately map RNA-seq reads onto the genome^77,78^. In brief, we used STAR v2.4.2a^79^ to align RNA-seq reads to the reference (parameters: -- outFilterMultimapNmax 20 --outFilterMismatchNmax 999 --outFilterMismatchNoverReadLmax 0.07 -- alignIntronMin 20 –alignIntronMax 1000000 --alignMatesGapMax 1000000 --alignSJoverhangMin 8). The reference genomes used were: human, hg19 and mouse, mm9. Gene models were obtained through the UCSC Genome Browser for Gencode, RefSeq, Ensembl, and UCSC Genes. We only considered uniquely mapped paired-end reads with mapping quality q=255 and used picard-tools (http://broadinstitute.github.io/picard) to remove PCR duplicates mapped to the exact same location. Uniquely mapped reads were subjected to local realignment and base score recalibration using the IndelRealigner and TableRecalibration tools from the Genome Analysis Toolkit (GATK)^80^.

### Gene expression analysis

The expression of known genes (measured in Transcripts Per Kilobase Million, TPM) was quantified using RSEM v1.2.21^81^ on the basis of STAR mappings for all RNA-seq libraries (parameters: --bam -- estimate-rspd --calc-ci --seed 12345 --paired-end --forward-prob 0). Gene models were obtained from Ensembl for human (Release 72) and mouse (Release 67). If one editing site overlapped with models of genes transcribed in the opposite directions, we used the strand information from RNA-seq to calculate strand-specific read count for downstream analysis. List of ISGs (Interferon-Stimulated Genes) is provided in **Table S6**.

### De novo identification of A-to-I RNA editing site

De novo identification of editing sites was performed on pooled mapping results of HEK293T and NPC samples. We used the UnifiedGenotyper tool from the Genome Analysis Toolkit (GATK)^80^ to call variants from mapped RNA-seq reads. We then applied parameters and filters as described previously^77,78^. In brief, human variants at non-repetitive and repetitive non-*Alu* sites were required to be supported by at least three mismatch reads. A support of one mismatch read was required for variants in human *Alu* regions. This set of variant candidates was subject to several filtering steps that increased the accuracy of editing site discovery^77,78^. We also removed all known SNPs present in dbSNP (except SNPs of molecular type “cDNA”; database version 135; http://www.ncbi.nlm.nih.gov/SNP/), the 1000 Genomes Project, and the University of Washington Exome Sequencing Project (http://evs.gs.washington.edu/EVS/). We also performed hyper-editing identification to recover the editing sites from unmapped reads^82^. At last, we combined known editing sites from RADAR and the newly identified sites from each cell line, and obtained 3,949,224 and 2,982,889 editing sites in HEK293T and NPC, respectively.

### Identification of A-to-I RNA editing clusters

To identify the long dsRNAs that are potential MDA5 ligands without relying on any prior annotation, we first reasoned that these long dsRNAs should all be hyper-edited in order to suppress their immunogenicity. We took the location information of editing sites in the genome as proxy for dsRNAs and applied a density-based clustering algorithm (DBSCAN)^83^ to identify genomic regions with many editing sites close together, which we called editing clusters. DBSCAN requires two parameters to define a cluster: the minimum number of points (editing sites) required to form a cluster (minPts) and the farthest distance between two adjacent sites within the same cluster (eps). For the first parameter (minPts), as expected, we observed that the total number of clusters decreases as the minimum number of editing sites per cluster increases (**Extended Data Fig. 3b**). Based on *in vitro* MDA5 binding data from this and previous studies^6^, we set minPts=5 as the minimum number of mismatches (edits). For the second parameter (eps), by plotting the distance between adjacent sites sorted from the nearest to the farthest, we observed a “knee” point on the curve, where eps = 120 (**Extended Data Fig. 3c**). Since this “knee” point allows the inclusion of >90% editing sites in editing clusters while excluding the far away outliers, we concluded that eps = 120 is the optimal value. By applying DBSCAN clustering, we identified 205,652 and 175,655 editing clusters in HEK293T and NPC cell lines, respectively. Further after applying the coverage cutoff (≥20 reads), we obtained 39,079 and 40,182 editing clusters in HEK293T and NPC cell lines for downstream analyses, respectively.

### Quantification and comparison of editing index

We adopted the concept of an “editing index” as the measurement of the editing level of an editing cluster. For each cluster containing *n* editing sites, its editing index is calculated as: 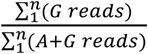. To ensure the measurement of editing index is reliable, we required that: 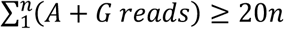. For each editing cluster meeting the above threshold in both conditions in comparison, Fisher’s exact test was applied to the counts of G and A reads to calculate the p-value for the significance of the editing indices not being equal. Multi-test p-values were adjusted using Benjamini–Hochberg procedure^84^.

### Neural cell type enrichment analysis

In order to identify the relevant neural cell types in which the immunogenic dsRNAs are most likely to be expressed, we performed expression-weighted celltype enrichment (EWCE) analysis^58^. First, we calculated expression specificity matrices using public single-cell transcriptome data of mouse cortex and hippocampus^85^. Then we took the lists of genes containing immunogenic dsRNAs identified specifically in NPC and HEK293T cells, and kept only the ones with mouse orthologs. An enrichment test was then performed on the mouse orthologs, controlling for transcript length and GC content, along with 10,000 bootstrap lists to determine significance. Lastly, we reported multiple test corrected p-values and cell-type enrichment scores (calculated as standard deviations from the mean).

### Characterization of dsRNA features

For dsRNAs formed by IR*Alu*s, we used the RNAfold function from the ViennaRNA package^86^ to calculate minimum free energy (mfe) secondary structures (default parameters). We predicted the mfe secondary structures using the closest inverted *Alu* pairs plus the non-*Alu* sequences in between the members of the pairs. We also computed the partition function to obtain the frequency of the mfe structure in the ensemble, as well as the ensemble diversity. Based on the predicted mfe structure, we calculated the fraction of base pairs as well as the number of mismatches/bulges. All quantitative dsRNA features were normalized and analyzed using principal component analysis (PCA) to compute each feature’s relative contribution.

### Enrichment analysis of dsRNAs in GWAS signal

To make Q-Q plots of GWAS signal annotated with edQTL associated with dsRNAs, we clumped the significant edQTLs to obtain independent signals using PLINK^87^ (--clump-r2 0.4 –clump-kb 250). To control for differences in the total number between immunogenic dsRNAs and non-immunogenic dsRNAs, we randomly selected non-immunogenic dsRNAs of equal number as the immunogenic dsRNAs, which also had edQTL in the GTEx edQTL^13^. The summary statistics of GWAS were obtained as described in our recent work^13^.

### Quantification and statistical analysis

For qRT-PCR, mRNA expression was examined using the ddCt method with Actin as the internal normalization control. Statistical analyses were performed in Graph Pad PRISM 8. Comparisons between groups were made using the Student’s *t*-test.

### Data and code availability

The accession number for the sequencing data reported in this paper is GEO: GSE155964. A secure token (izwlcogqlnsxtmp) has been generated to allow review of GSE155964 while it remains in private status. Codes used in this study are available from the corresponding author on reasonable request.

## Acknowledgments

We thank Prof. Carl Walkley (St Vincent’s Institute of Medical Research) for the MDA5 constructs, helpful discussion and critical reading of the manuscript. We thank Prof. Charles Rice and Dr. Hachung Chung (Rockefeller University) for sharing human ADAR1 mutant cells and the relevant protocol. We thank Prof. Lei Stanley Qi (Stanford University) for sharing the RfxCas13d plasmid. We acknowledge Dr. Xin Liu and Ms. Sandra Linder for technical support.

## Author Contributions

T.S., Q.L. and J.B.L. conceived of the project. T.S. performed most of the experiments with help from J.M.G., S.-B.H., B.F., S.S., H.G, J.M. and J.B.L. Q.L. performed computational analyses with the help of J.B.L. The manuscript was written by T.S., Q.L. and J.B.L. with input from other authors.

## Declaration of Interests

J.B.L. is a co-founder of AIRNA Bio and a consultant for Risen Pharma.

Correspondence and requests for materials should be addressed to.

**Supplementary Information** is available for this paper.

**Tables S1**.

Putative immunogenic dsRNAs identified in HEK293T cells.

**Tables S2**.

Putative immunogenic *cis*-NATs in HEK293T and NPCs.

**Tables S3**.

Putative immunogenic dsRNAs identified in NPCs.

**Tables S4**.

Gene expression of components in MDA5 and type-I IFN pathways.

**Tables S5**.

Sequences of DNA oligonucleotides.

**Tables S6**.

Interferon-Stimulated Genes (ISGs) used in this study.

