## Supplemental Figures 1-10 for "A Small Subset of Cytosolic dsRNAs Must Be Edited by ADAR1 to Evade MDA5-Mediated Autoimmunity"

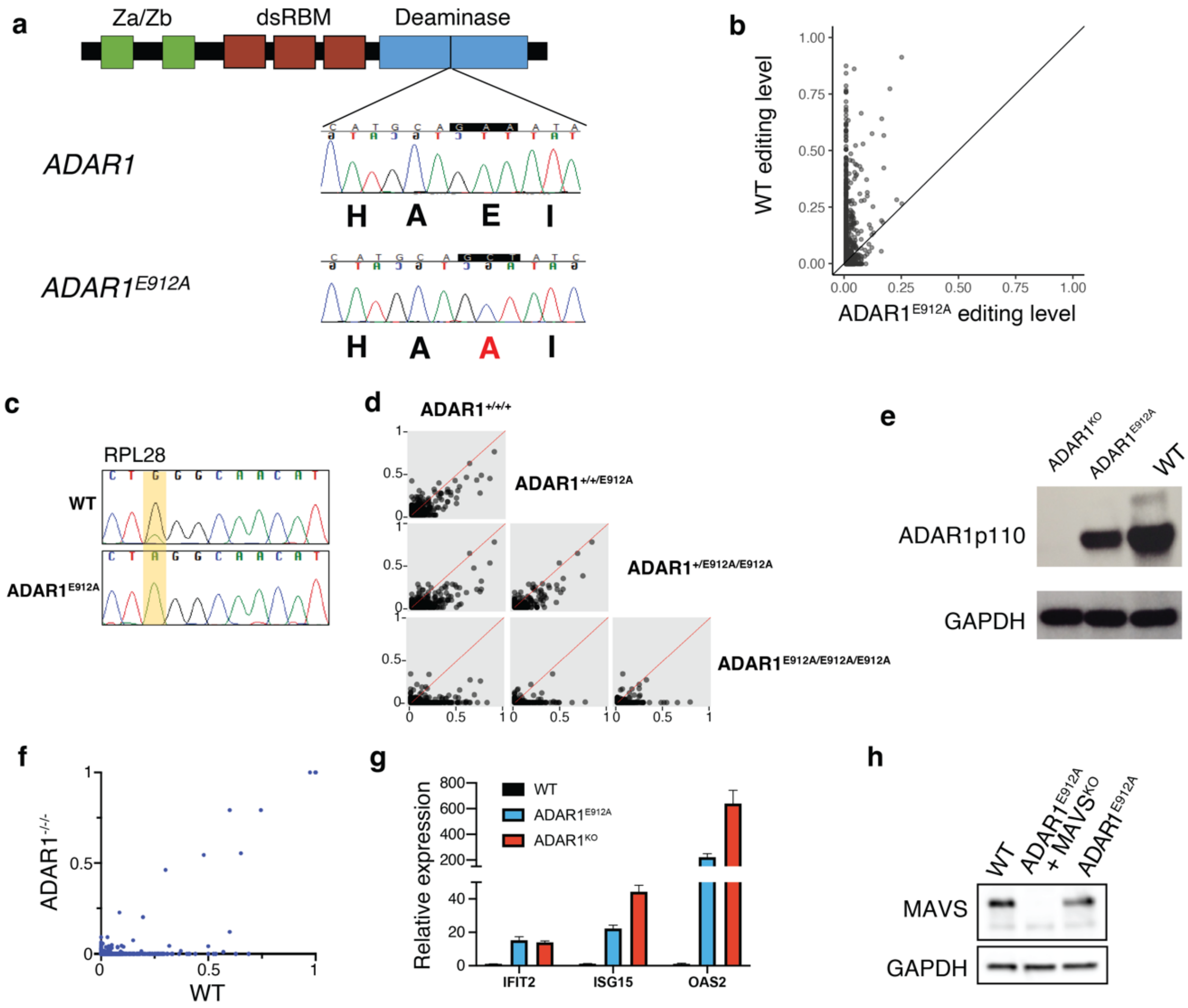


**Extended Data Figure 1: Generation and characterization of ADAR1$^{E912A}$ and ADAR1$^{KO}$ mutants in HEK293T cells.** **a**, A homozygous *ADAR1$^{E912A}$* mutant is generated in HEK293T cells using CRISPR knock-in. The chromatogram from Sanger sequencing of the targeted region indicates all alleles are mutated. The mutant has three alleles of *ADAR1*, and is denoted *ADAR1$^{E912A}$* for short. **b**, RNA editing level comparison between wild type (WT) and ADAR1$^{E912A}$ cells. RNA editing levels were quantified from total RNA-seq. Sites with ≥20 reads coverage were plotted. **c**, RNA editing level of an editing site (highlighted in yellow) in RPL28 is shown in the chromatogram of Sanger sequencing in WT and ADAR1$^{E912A}$ cells. **d**, Pairwise comparison of RNA editing levels between ADAR1$^{+/+/+}$ (WT), ADAR1$^{+/+/E912A}$, ADAR1$^{+/E912A/E912A}$ and ADAR1$^{E912A/E912A/E912A}$ (hereafter ADAR1$^{E912A}$). **e**, Western blot of ADAR1 and GAPDH proteins in ADAR1$^{KO}$ (ADAR1$^{-/-/-}$), ADAR1$^{E912A}$ and ADAR$^{WT}$ cells. **f**, RNA editing level between WT and ADAR1$^{-/-/-}$ cells. **g**, Relative expression levels of three representative ISGs (IFIT2, ISG15, and OAS2) in WT, ADAR1$^{E912A}$ and ADAR1$^{KO}$ cells with MDA5 expressed. Data are normalized to WT expression level and represented as mean ± SD (technical replicates n = 4). **h**, Western blot of MAVS and GAPDH proteins in WT, ADAR1$^{E912A}$ + MAVS$^{KO}$ and ADAR1$^{E912A}$ cells.

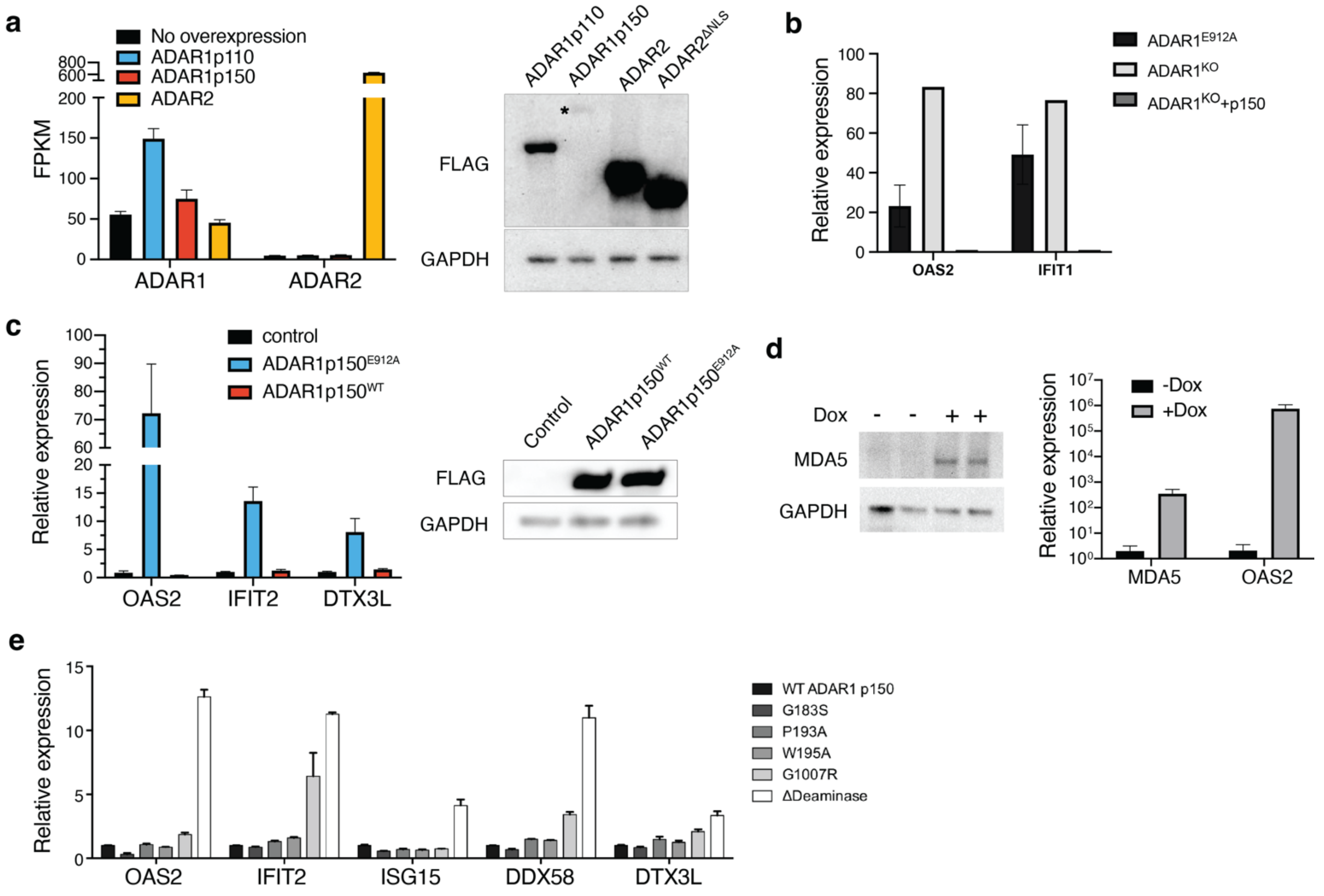


**Extended Data Figure 2: ADAR protein isoforms and functional domains. a**, Expression level of ADAR1 or ADAR2 in different cells lines measured by RNA-seq (left) and western blot (right). Black bar represents ADAR1$^{E912A}$ cell line and each colored bar represents an engineered ADAR1$^{E912A}$ cell line with an ADAR isoform overexpressed. For western blot, Flag-tagged ADAR1 and ADAR2 isoforms were stably expressed in the ADAR1$^{E912A}$ cells via lentivirus transduction. Expression was detected by western blot using the FLAG antibody. GAPDH was used as an internal control. ADAR1p150 expression is relatively low and marked with *. **b**, Relative expression levels of two ISGs (OAS2 and IFIT1) in ADAR1$^{E912A}$, ADAR1KO and ADAR1KO+p150 cells with MDA5 expressed (technical replicates n=2). **c**, Left panel, relative expression levels of three ISGs (OAS2, IFIT2 and DTX3L) in WT cells and ADAR1$^{E912A}$ cells expressed with ADAR1p150$^{E912A}$ and ADAR1p150$^{WT}$ (technical replicates n=3). Right panel, western blot of FLAG-tagged ADAR1p150$^{E912A}$ and ADAR1p150$^{WT}$ proteins. Expression was detected by western blot using the FLAG antibody. GAPDH was used as an internal control. **d**, ISG induction in ADAR1$^{E912A}$ HEK293T cells with Dox-inducible MDA5. Left, Western blot of MDA5 and GAPDH proteins without and with Dox induction. Right, RNA expression of MDA5 and OAS2 without and with Dox induction (biological replicates n=2). **e**, Relative expression levels of five ISGs (OAS2, IFIT2, ISG15, DDX58 and DTX3L) in ADAR1E912A cells expressed with WT ADAR1p150 and different mutants of ADAR1p150 including ΔDeaminase with the entire deaminase deleted (n=3). All data are represented as mean ± SD.

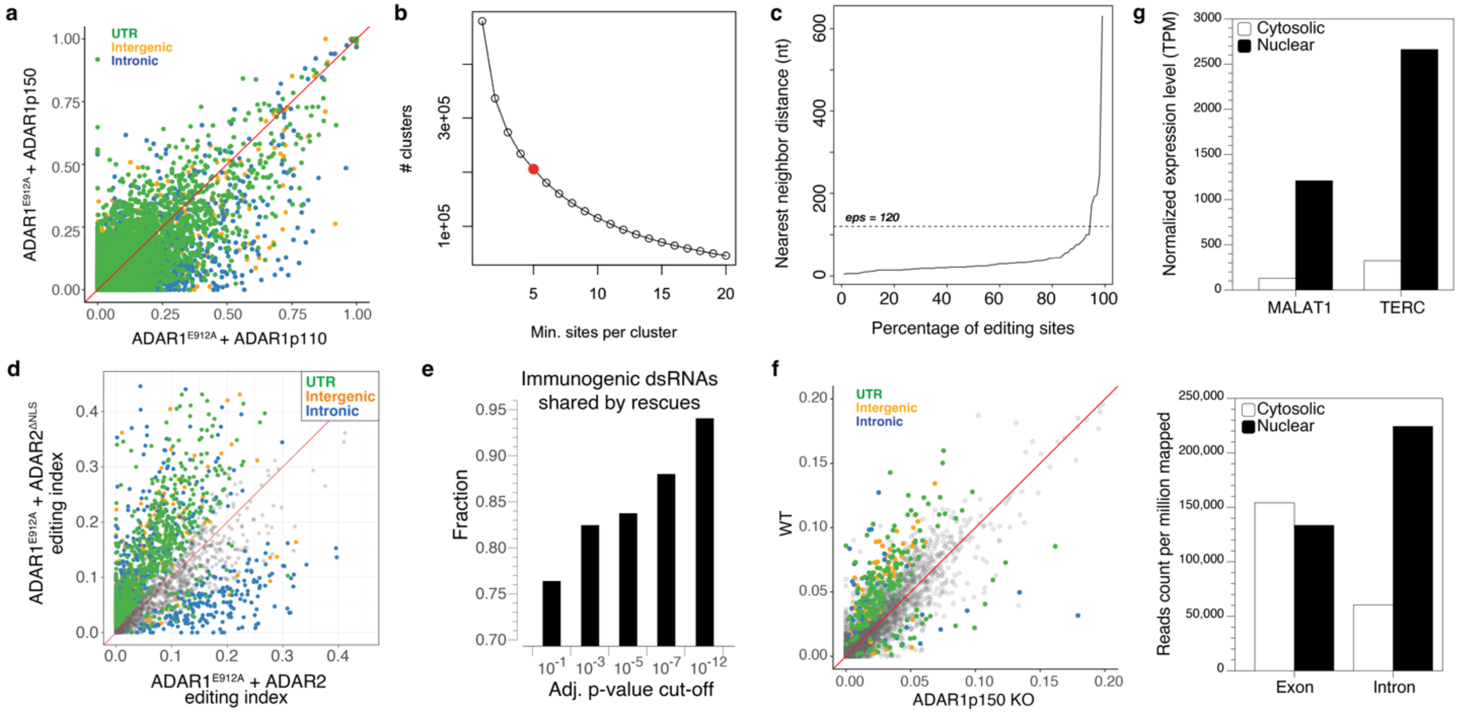


**Extended Data Figure 3: Editing status comparison. a**, Editing level of individual editing sites between ADAR1$^{E912A}$+ADAR1p110 and ADAR1$^{E912A}$+ADAR1p150. Sites with significantly different editing levels (adjusted p-value < 0.01) were colored based on genomic location (UTR, intergenic or intronic). **b**, Number of editing clusters in relation with the minimum sites per cluster. Red dot indicates the number of clusters when n = 5. **c**, Systematic method for determining the eps value. The distance between the nearest neighbor editing sites is plotted from the lowest to the highest. The knee point of the curve is determined using kneed (https://github.com/arvkevi/kneed). **d**, Editing index comparison between ADAR1$^{E912A}$ cells complemented with ADAR2 and ADAR2$^{\Delta NLS}$ The editing clusters with significant index differences are color coded the same as in **a**. Nonsignificant clusters are shown in grey. **e**, The fraction of immunogenic dsRNAs identified from the ADAR1p150 vs. ADAR1p110 comparison that were also identified from the ADAR2 vs. ADAR2$^{\Delta NLS}$ comparison. The cut-off of adjusted p-values of multiple tests following Benjamini-Hochberg correction (FDR = 0.05) applied to both ADAR1p150 vs ADAR1p110 and ADAR2 vs. ADAR2$^{\Delta NLS}$ comparisons. **f**, Editing index of dsRNA clusters between WT and ADAR1p150 KO cells. The editing clusters with significant index differences are color coded the same as in **a**. Nonsignificant clusters are shown in grey. **g**, RNA-seq analysis of cytosolic and nuclear fractionations. Top panel, expression level of nuclear marker genes *MALAT1* and *TERC* in cytosolic and nuclear fractionation samples. Expression level is normalized to TPM (Transcripts Per Kilobase Million). Bottom panel, total number of reads mapped to exons and introns in cytosolic and nuclear fractionation samples. The total mapped reads count is normalized to one million reads per sample.

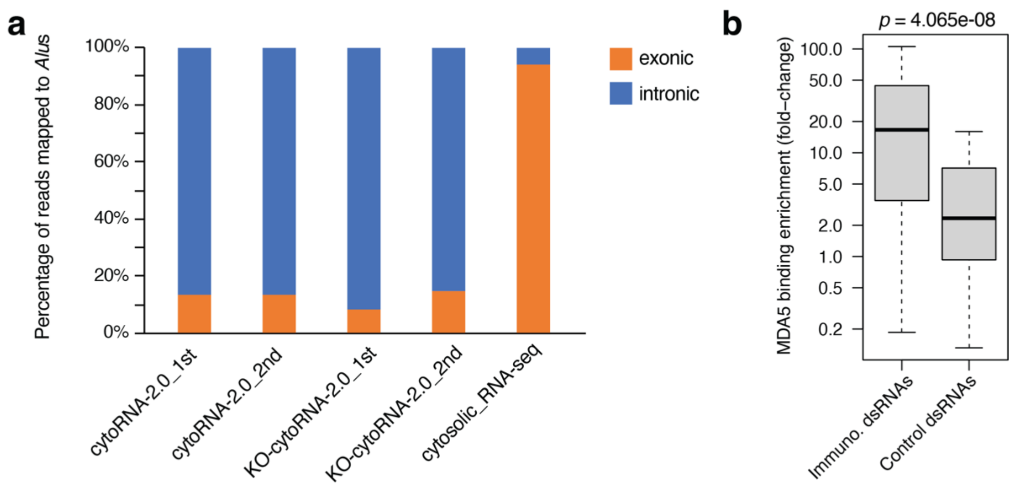


**Extended Data Figure 4: Analysis of the RNA identified from RNase protection assay. a**, Percentage of RNA sequencing reads mapped to *Alu*s in exonic and intronic regions from the RNase protein assay experiments and the control of cytosolic RNA-seq. cytoRNA-2.0_1st and cytoRNA-2.0_2nd: two replicates of RNase protection assay using cytosolic RNA of WT HEK293T and Gain-of-Function MDA5 protein. KO-cytoRNA-2.0_1st and KO-cytoRNA-2.0_2nd: two replicates of RNase protection assay using cytosolic RNA of ADAR1 KO HEK293T cells and WT MDA5 protein. cytosolic_RNA-seq: RNA-seq of total cytosolic RNA from WT HEK293 cells. **b**, Enrichment of MDA5 binding to dsRNAs in ADAR1 KO cells. MDA5 binding enrichment was quantified by the normalized expression fold-change of dsRNAs in RNase-treated samples compared to untreated samples. Control dsRNAs were randomly selected non-immunogenic dsRNAs located in 3' UTR with comparable expression to the immunogenic dsRNAs with matched numbers. Median fold-change of immune. dsRNAs vs. control dsRNAs: 16.6±6.3 vs. 2.34±2.08, *p*-value = 4.065e-08, Wilcoxon signed-rank test. The sequencing data for the RNase protection assay were obtained from Ahmad et al., 2018 (GEO: GSE104865).

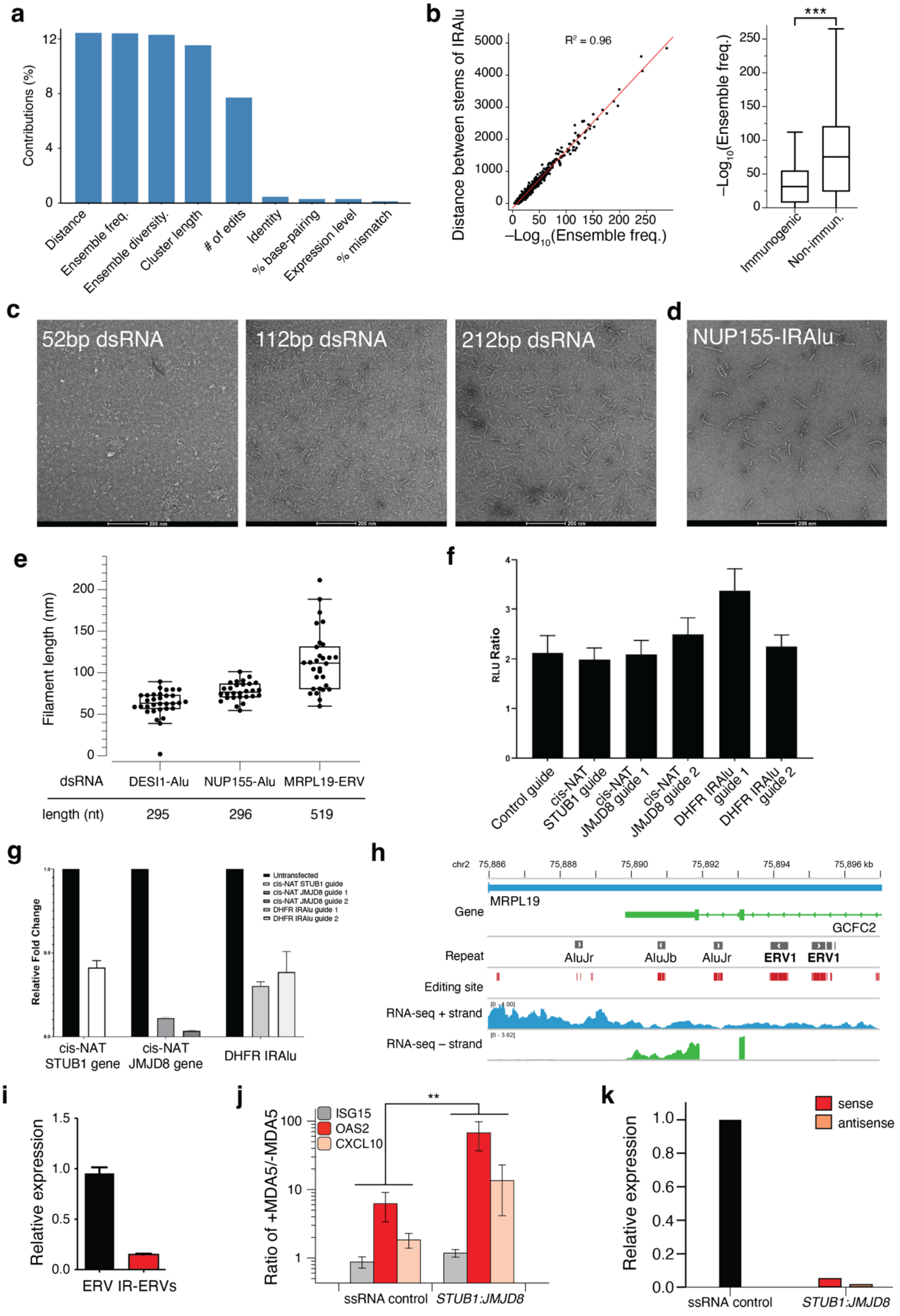


**Extended Data Figure 5: Characterization of immunogenic dsRNAs. a**, The contribution of IR*Alu* features to the immunogenicity in PCA. **b**, Comparison between ensemble frequency of dsRNA with the distance between IR*Alu*s (left). Comparison of ensemble frequency between putative immunogenic dsRNAs and non-immunogenic ones (right). *** p<0.001. **c**, Visualization of filament formed by MDA5 and dsRNA of different lengths using negative staining EM. **d**, Visualization of filament formed by MDA5 and IR*Alu* dsRNAs located in *NUP155* 3'UTR. **e**, Filament length comparison of dsRNAs formed by inverted repeats. Lengths of the dsRNAs are indicated at the bottom of the table. Data are represented as interquartile range. **f**, The relative ratios of luminescence units in doxycycline-treated vs doxycycline-untreated cells that were transfected with plasmids bearing Cas13d and the indicated guide RNAs targeting the *STUB1:JMJD8 cis*-NAT or IR*Alu* in *DHFR*. **g**, qPCR to measure knockdown efficiencies of guide RNAs depicted in **f**. RNA from transfected HEK293T cells were harvested 48 hours after transfection. Data graphed as mean +/- SEM, $n$ = 3. **h**, Genome browser view of the IR-ERVs and editing sites embedded in the 3'UTR of *MRPL19*. IR-ERVs are transcribed as part of MRPL19 3'UTR as supported by the stranded RNA-seq data. **i**, Real-time PCR measurement of overexpression of ERV or IR-ERV derived from *MRPL19*. Plasmid with single or inverted repeat (IR) ERVs at the 3'UTR of a GFP was transfected into the cells. Real-time PCR was performed on the co-expressed fluorescent proteins GFP to quantify the expression level of the ERV. The expression was normalized to the internal control Actin for ERV. Biological replicates n=2, technical replicates n=3. **j**, Ratio of expression of three signature ISGs for cells in the presence of doxycycline-induced MDA5 versus its absence. Cells were transfected with a plasmid coexpressing the *STUB1:JMJD8 cis*-NAT pair or the pKER-mClover3 ssRNA control. Biological replicates n>=3. **: $p$ < 0.01, One-way ANOVA test. **k**, Real-time PCR measurement of overexpression of *STUB1:JMJD8 cis*-NAT relative to mClover3 ssRNA control. Overexpression of sense strand (*STUB1*) was measured by the co-expressed mClover3 and antisense strand (*JMJD8*) by mRuby3.

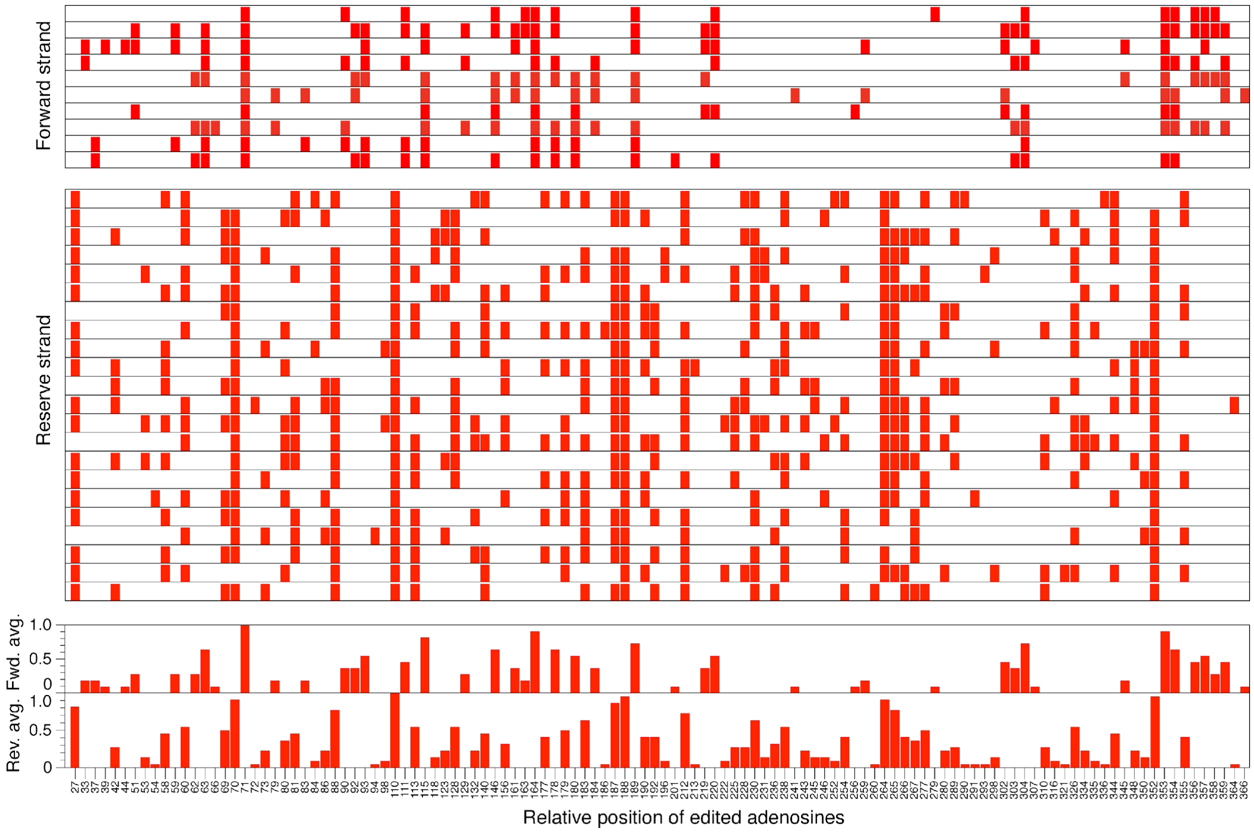


**Extended Data Figure 6: *In vitro* editing status of 32 TOPO clones of dsRNAs formed by *cis*-NATs of *JMJD8:STUB1*.** Rows represent individual clones while columns represent editable adenosines in the overlapping *cis*-NATs. Both forward and reverse strands of *cis*-NATs can be edited and the editing status is shown separately for the two strands. Average editing level of each adenosine position across clones is shown at the bottom. Editing information was measured using Sanger sequencing.

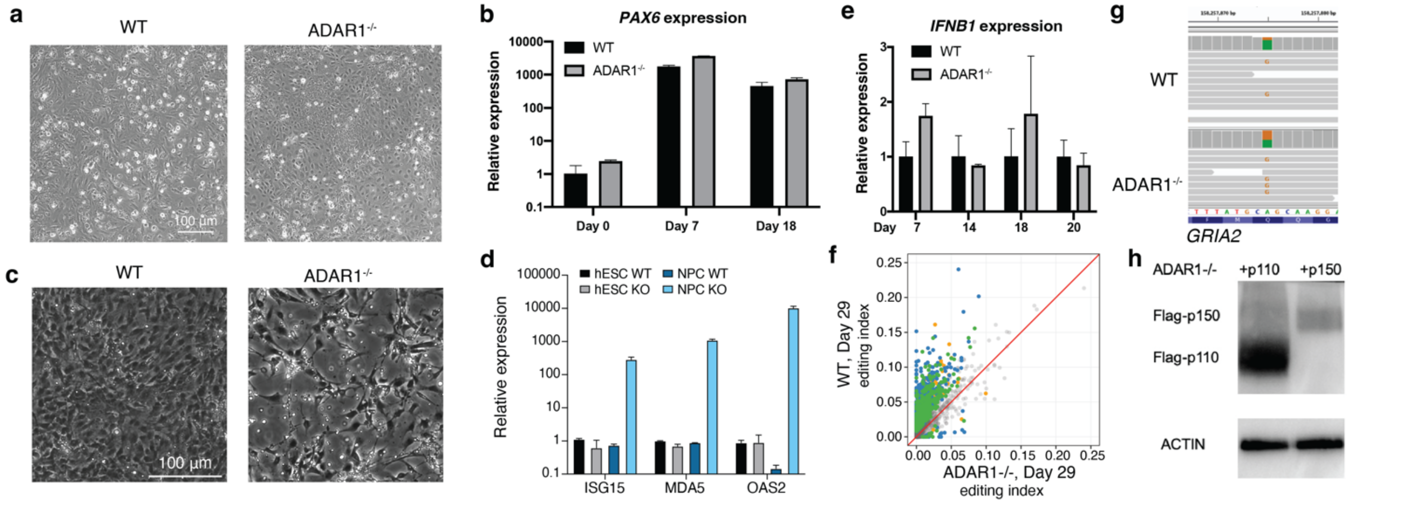


**Extended Data Figure 7: Characterization of the role of ADAR1 in NPCs. a**, Cell morphology of the differentiated NPC 7 days post induction. **b**, NPC marker PAX6 expression at different time points post induction in WT and ADAR1$^{-/-}$ cells (Biological replicates n=3). **c**, Cell morphology of the WT and ADAR1$^{-/-}$ NPCs at Day 29 after differentiation. **d**, Signature ISG expression in hESC and NPC at Day 29 after differentiation. Real-time PCR was performed on RNA of WT and KO (ADAR1$^{-/-}$) cells (Biological replicates n=2). **e**, RNA expression of IFNB1 at different time points post induction in WT and ADAR1$^{-/-}$ NPCs (Biological replicates n=2). **f**, Editing index comparison between ADAR1p110- and ADAR1p150-expressing ADAR1$^{-/-}$ NPCs. Editing clusters with significantly different editing index (adjusted p-value < 0.01) were colored the same way as in **Extended Data Fig. 3a**. **g**, RNA editing level of ADAR2-specific GRIA2 Q/R site in the WT and ADAR1$^{-/-}$ NPCs. RNA-seq track is visualized by IGV and the edited base ‘G’ is highlighted. Data are represented as mean ± SD for **b**, **d** and **e**. **h**, Protein level of ADAR1p110 and ADAR1p150 in rescued ADAR1$^{-/-}$ NPCs. ADAR1$^{-/-}$ NPCs were transduced with lentivirus to over-express Flag-tagged ADAR1p110 or ADAR1p150 isoform. The cells were selected by puromycin before the total protein was assayed by western blot using the anti-Flag antibody. Actin was used as an internal loading control.

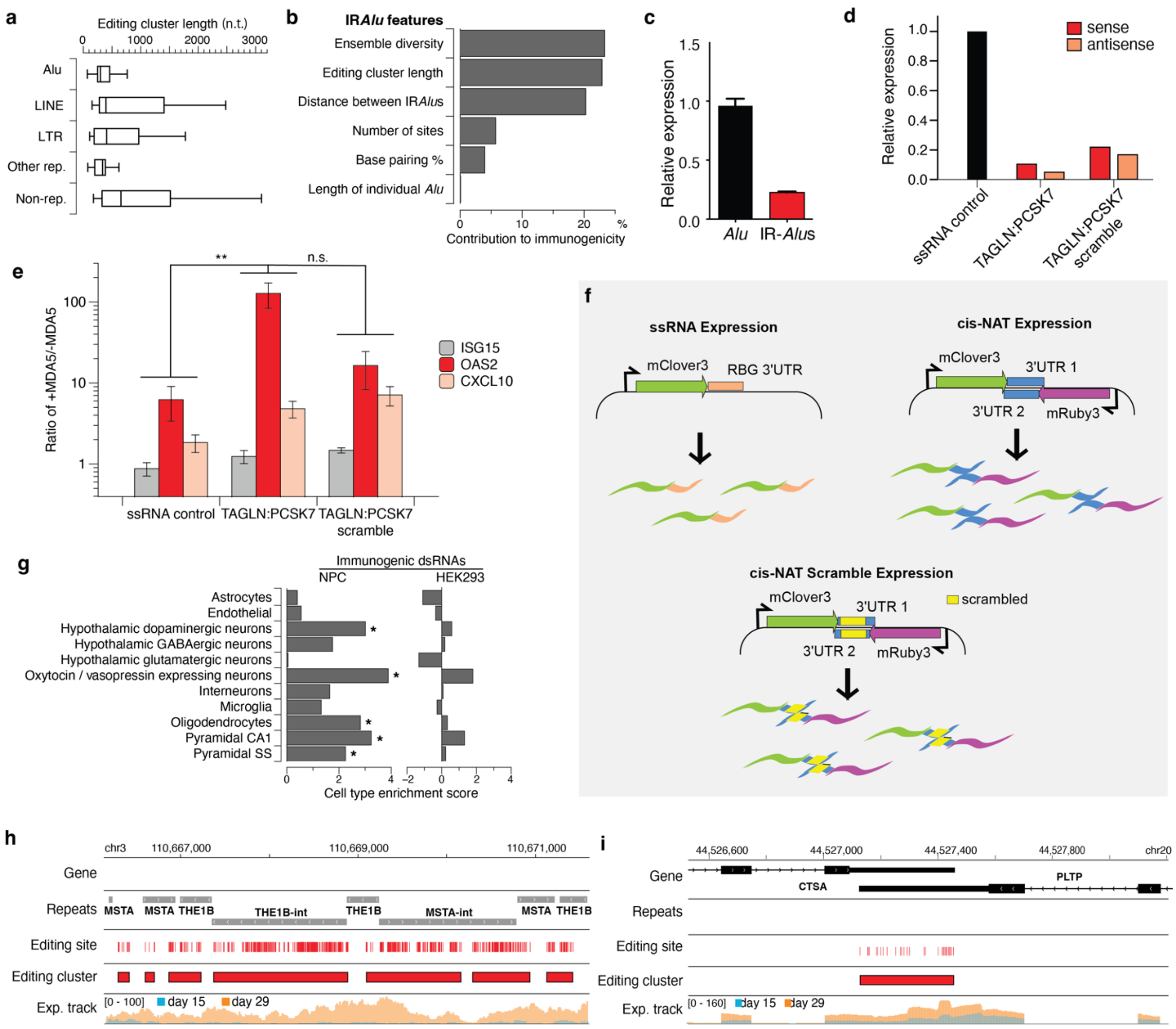

**Extended Data Figure 8: Characterization of immunogenic dsRNAs in NPCs. a**, Editing cluster length comparison among different classes of immunogenic RNAs. Data were represented as interquartile ranges. **b**, Contribution of different IR*Alu* features to dsRNA immunogenicity. **c**, Real-time PCR measurement of expression of *Alu* or IR-*Alu* derived from *NICN1*. Plasmid with single or inverted repeat (IR) *Alu*s at the 3'UTR of a GFP was transfected into the cells. Real-time PCR was performed on the co-expressed fluorescent proteins GFP to quantify the expression level of the *Alu*. The expression was normalized to the internal control Actin for *Alu*. Biological replicates n=2. **d**, Relative expression level of *TAGLN:PCSK7 cis*-NAT to mClover3 ssRNA control overexpression determined via RT-qPCR. Overexpression of sense strand (*TAGLN*) was measured by the co-expressed mClover3 and antisense strand (*PCSK7*) by mRuby3. **e**, Ratio of expression of selected ISGs in the presence vs absence of doxycycline-induced MDA5 from cells transfected with a plasmid expressing the *cis*-NAT pair *TAGLN:PCSK7* or with the pair's overlapping sequence scrambled. Data depicted as mean ± SEM. Biological replicates n >= 3. n.s.: $p \geq 0.05$, **: $p < 0.01$, One-way ANOVA test. **f**, Schematic depicting overexpression strategy for ssRNA control versus *cis*-NATs and *cis*-NATs containing scrambled sequence in place of the hyperedited overlapping region (indicated in yellow). **g**, Neuronal cell-type enrichment analysis of NPC- and HEK293-specific immunogenic dsRNAs. *, adjusted p<0.01. Pyramidal CA1, hippocampal CA1 pyramidal neurons. Pyramidal SS, hippocampal somatostatin pyramidal neurons. Examples of immunogenic dsRNA editing status and expression levels between Day 15 and Day 29 post-differentiation of NPCs are shown for ERVs in panel **h** and *cis*-NATs in panel **i**.

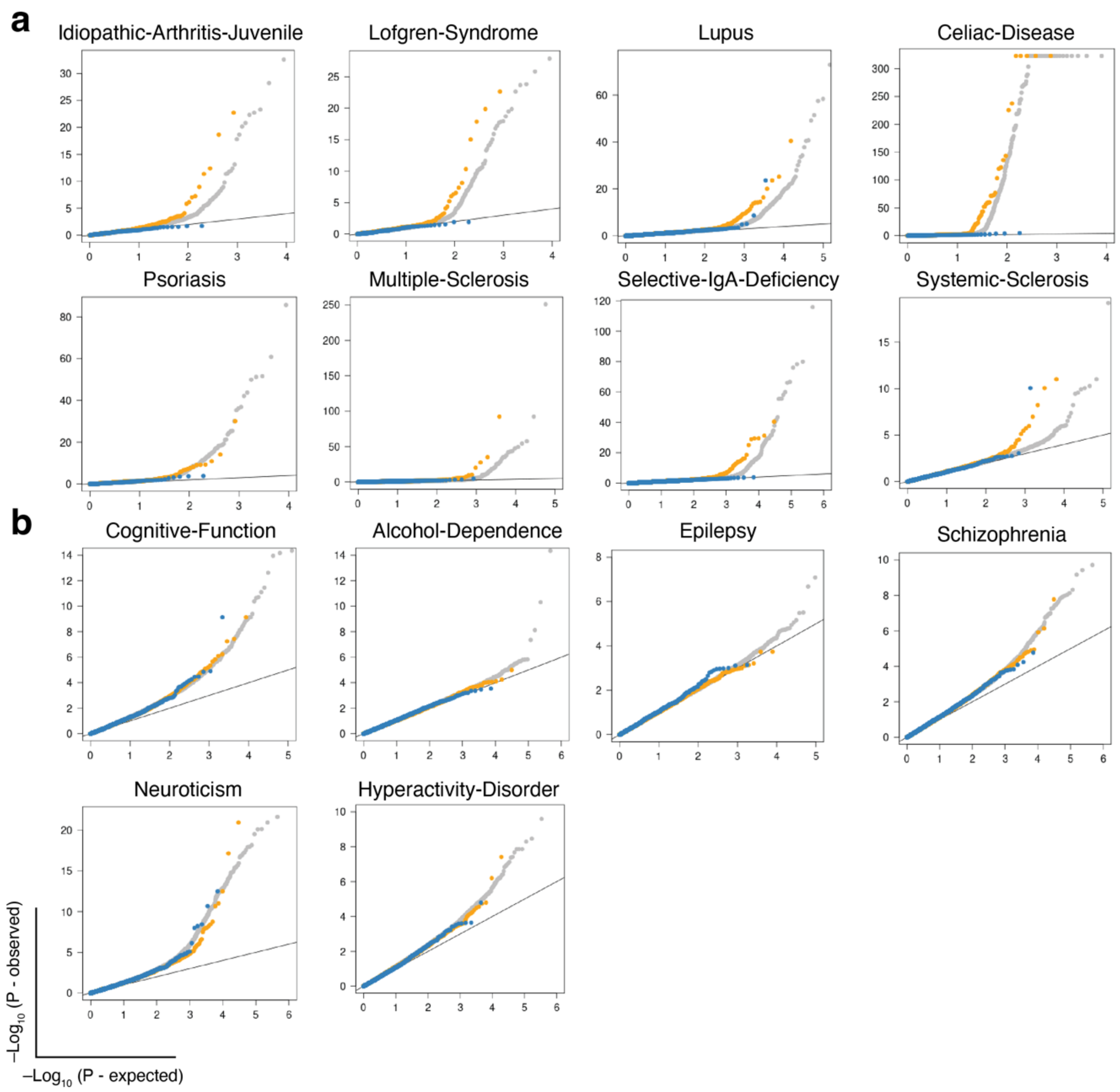


**Extended Data Figure 9: Comparison of immunogenic vs. non-immunogenic dsRNAs for enrichment in disease GWAS. a,** Quantile-Quantile (Q-Q) plots comparing enrichment for immunogenic dsRNAs (orange dots) vs. non-immunogenic dsRNAs (blue dots) in GWAS of 8 inflammatory diseases (grey dots). **b**, Quantile-Quantile (Q-Q) plots comparing enrichment for immunogenic dsRNAs (orange dots) vs. non-immunogenic dsRNAs (blue dots) in GWAS of 6 non-inflammatory diseases or traits (grey dots).

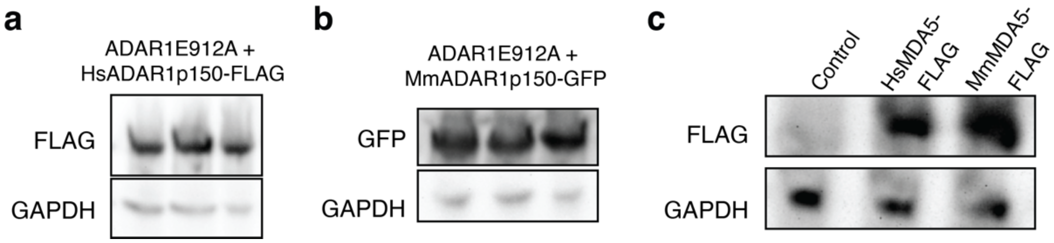


**Extended Data Figure 10: Expression of ADAR1 and MDA5 of mouse or human origin. a**, human and **b**, mouse ADAR1p150 overexpression in the human ADAR1$^{E912A}$ cells. Human ADAR1p150 was tagged with FLAG and mouse ADAR1p150 was tagged with GFP. They were detected with the FLAG or GFP antibody respectively. Technical replicates n=3. **c**, human and mouse MDA5-FLAG overexpression in the human ADAR1 KO cells. MDA5 was detected by the FLAG antibody.
